# Seasonal structural stability promoted by forest diversity and composition explains overyielding

**DOI:** 10.1101/2024.03.11.584423

**Authors:** J. Antonio Guzmán Q, Maria H. Park, Laura J. Williams, Jeannine Cavender-Bares

## Abstract

The stability of forest productivity over time is a widely studied parameter often associated with benefits of forest diversity. Yet, the structural stability (*SS*) through the season of forest communities and its relationship to diversity, composition, and productivity remains poorly understood. Using a large-scale (10 ha) young tree diversity experiment, we evaluated how forest structure and multiple dimensions of diversity and composition affect remotely-sensed structural metrics and their stability through the growing season. We then studied the impact of *SS* across the season (April-October) on the net biodiversity effects of annual wood productivity (i.e., overyielding) of forest mixtures. We surveyed experimental tree communities eight times at regular intervals from before bud-break to after leaf senescence, using an UAV-LiDAR to derive metrics associated with canopy height heterogeneity, gap probability, and forest structural complexity (i.e., fractal geometry). The inverse coefficient of variation of these metrics through the season was used as descriptors *SS*. These metrics along with their *SS* were then coupled with annual tree inventories to evaluate their relationships. Our findings indicate that plot wood volume and, to some extent, multiple dimensions of diversity and composition (i.e., taxonomic, phylogenetic, and functional) influence remotely-sensed metrics of forest structure and stability over time. We found that increases in plot wood volume as well as functional and phylogenetic diversity and variability (a measure of diversity independent of species richness), are linked to higher structural stability of forest complexity over time. We further found that higher stability of forest structural complexity and tree cover (i.e., 1 - gap probability) increases net biodiversity effects in forest mixtures through species complementarity. Structural equation models indicate that structural stability explains more the variation among plots in net biodiversity effects than multiple dimensions of diversity or variability, highlighting it as a measure that integrates several contributors to net biodiversity effects. Our results provide evidence that diversity and composition promote temporal stability of remotely-sensed forest structure and, in turn, enhanced productivity. The study highlights the potential to integrate remote sensing and ecology to disentangle the role of forest structural stability into ecological processes.

## Introduction

Diverse forest communities are critical for the maintenance of a healthy and resilient ecosystem (Messier et al. 2022). Diverse communities are known to enhance ecosystem functions, including nutrient cycling, soil microbial processes, and forest productivity. Among these benefits, forest productivity and its stability through time are the most studied parameters associated with forest diversity. Increases in diversity have consistently been shown to positively influence productivity and stability. A demonstrated cause of increased productivity is the complementarity of tree crowns as species with diverse architectures fill different portions of the available space, minimizing overlap among trees and reducing the amount of light that reaches the forest floor (Seidel et al. 2013, Williams et al. 2017, 2021, Juchheim et al. 2019). Forest stability — measured as the temporal variation of forest attributes through time — has been proposed to be an emergent property of diversity explained by the insurance hypothesis, where diverse communities provide more guarantees that some species will maintain forest function even if others fail (Yachi and Loreau 1999, Loreau et al. 2021). Although diversity and stability are essential to forest ecosystem function, they are difficult to monitor by traditional means. Thus, developing tools to remotely sense diversity and functional stability are critical to advancing our understanding for managing biodiversity on land (Cavender-Bares et al. 2022).

Recent evidence from tree biodiversity experiments indicates that increases in species richness lead to stable forest communities via species-asynchrony, temporal variation in growth among species, where interannual variation in species growth buffers community productivity against species-specific stress-related productivity declines (Schnabel et al. 2019, 2021). This phenomenon has also been shown in natural mixtures where species-asynchrony can counterbalance the negative effects of climate gradients on the stability of forest productivity (del Río et al. 2022). Although most of the current body of knowledge about forest stability comes from long-term effects of tree diversity, short-term effects (e.g., across seasons within a year) of diversity on stability have received less attention. Forest stability can be important in a single growing season given the variation in phenological development and growth patterns among species. From this perspective, species-asynchrony within a growing season may result from species-specific phenological rhythms (e.g., differences in the timing of leaf flushing or senescence) and responses to environmental fluctuations that are likely to influence forest structure (Smith et al. 2019). Consequently, diverse forest communities that integrate different growth and resource acquisition strategies across time are anticipated to sustain high levels of structural stability throughout the growing season. Structural stability, in turn, may help them maintain high levels of productivity. However, seasonal fluctuations in forest structure are difficult to assess using traditional forest inventory methods, which do not tend to capture forest structure in three dimensions.

Recent advances in remote sensing technologies are advancing our ability to frequently measure and monitor forest structure, diversity and composition (Turner 2014). Specifically, advances in Light Detecting and Ranging (LiDAR) are revolutionizing how we measure forest structure and structural diversity (Atkins et al. 2023a). Forest taxonomic diversity, particularly species richness, has long been associated with LiDAR structural metrics such as canopy height (Robinson et al. 2018), canopy height heterogeneity (Torresani et al. 2020), and structural complexity (Ehbrecht et al. 2017). Structural metrics derived from LiDAR metrics have been used to evaluate mechanisms associated with the net biodiversity effects (NBE) on productivity (i.e., overyielding) such as crown complementary (Kunz et al. 2019) and forest complexity (Ray et al. 2023). Despite this, it remains unclear how structural attributes of forests measured by LiDAR can be coupled to multiple dimensions of diversity and composition. Moreover, the assessment of structural stability across the growing season and its association with forest diversity, composition, and productivity is not well understood.

Here we aim to assess how multiple dimensions of forest diversity and composition are linked to forest structural metrics derived from LiDAR onboard to an Uncrewed Aerial Vehicle (UAV), and how forest structural variation throughout the growing season (i.e., structural stability, *SS*), which are influenced by diversity and composition, may contribute to net biodiversity effects. We focus our attention on three forest structural metrics derived from LiDAR, LiDAR-derived metrics herein: canopy height heterogeneity (*CH*_CV_) as a descriptor of the height variation hypothesis (Torresani et al. 2020, 2023), gap fraction probability (*P*_gap_) as a descriptor of light interception by forest openness, and the fractal dimension (*d*_D_) as metrics that captures the tridimensional arrangements of plant structures (i.e., forest structural complexity) (Guzmán et al. 2020). The variability of these metrics throughout the growing season was estimated as descriptors of structural stability. We took advantage of the low-density *Forest and Biodiversity* (FAB2) experiment (Cavender-Bares et al. in review) to address our goals. FAB2 provides an experimental platform from which to evaluate diversity effects with consistent forest age, tree density, climate, and management. We first hypothesize that increases in the volume occupied by a forest assemblage, measured on the ground as plot wood volume, strongly influence LiDAR-derived metrics and their structural stability. Second, we hypothesize that taxonomic, phylogenetic, and functional measures of diversity (which covary with species richness) and variability (which are independent of species richness), are likely to influence LiDAR-derived metrics of forest structure and stability (Figure 1a and b). Moreover, we hypothesize that communities composed of dissimilar species (i.e., high variability) are more likely to influence LiDAR-derived metrics of structural complexity and stability than diversity metrics that increase with species richness because assemblages with dissimilar species are prone to include a diverse range of tree architectures, phenological rhythms, and resource capture strategies. Finally, we hypothesize that diversity and variability contribute to structural stability resulting in increases in NBE (Figure 1c) because structural stability is an integrated measure that captures the emergent properties of interactions among species within communities. We test these hypotheses individually and examine their relative importance using structural equations models (Figure 1d).

**Figure 1.**
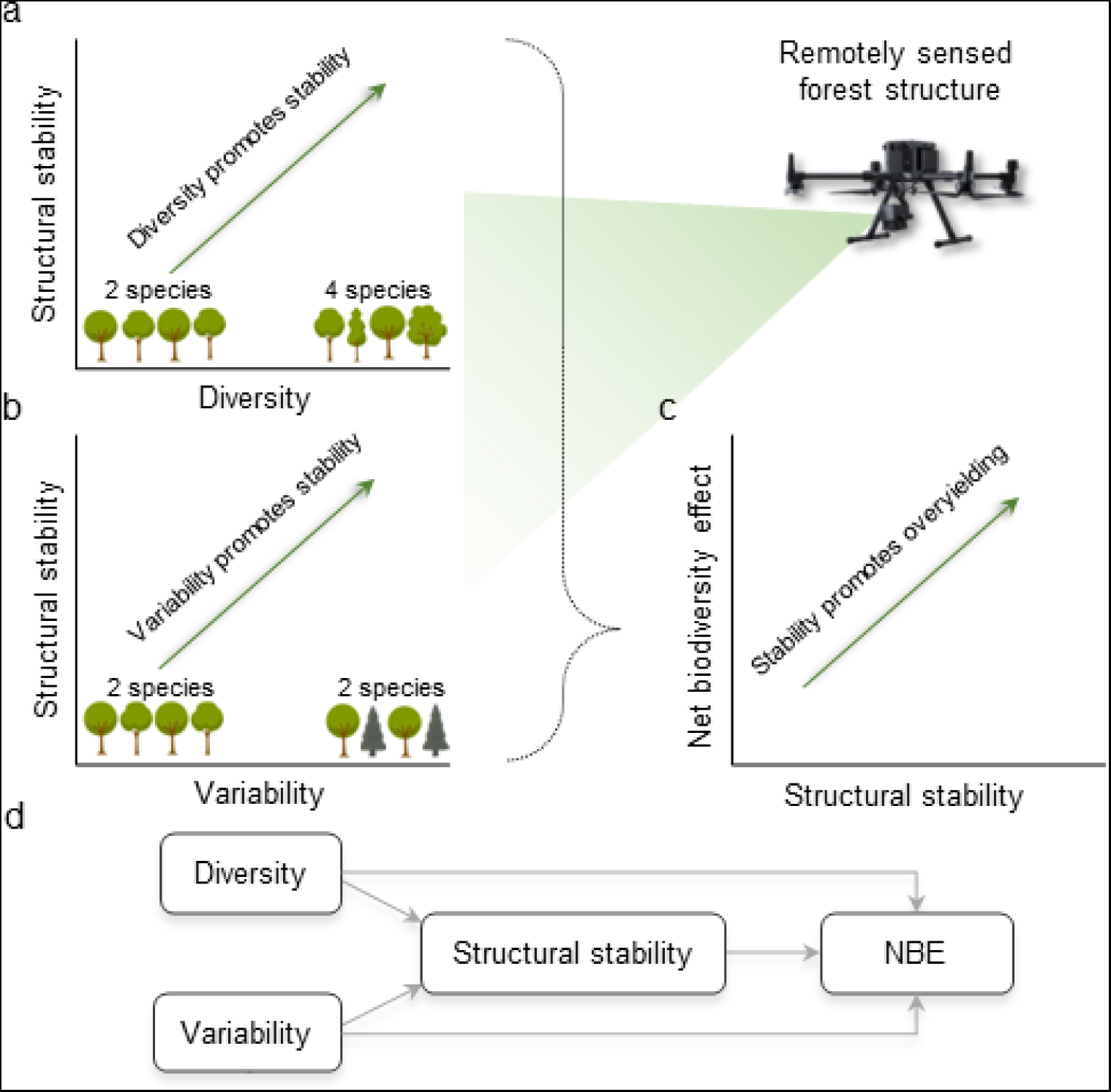
Schematic representation of how remotely-sensed structural stability through the growing season is promoted by multiple dimensions (i.e., taxonomic, phylogenetic, or functional) of diversity and variability which, in turn, affects net biodiversity (NBE) effects of productivity (i.e., overyielding). Panel ‘a’ describes expected trends with increases in diversity where trees resample species. Panel ‘b’ describes expected trends with variability (i.g., relatedness or similarity among species) where contrasting tree shapes and colors increases the community variability regardless of the number of species. Panel ‘c’ describes the expected trends of NBE indirectly promoted by diversity and variability. Panel ‘d’ integrates and compares our hypothesis using structural equation models.

## Methods and Materials

### Study site

We conducted this study in the Forest Biodiversity Experiment 2 (FAB2) located at the Cedar Creek Ecosystem Science Reserve (CCESR), Minnesota, USA (Cavender-Bares et al. in review) (45° 24’ 22.6" North and 93° 11’ 30.0" West). This site is characterized by a flat outwash plain with infertile, excessively drained soils consisting of upwards of 90% sand as part of the Anoka Sand Plain. This reserve has a humid continental climate with warm summers and cold winters. The region presents a mean annual temperature close to 7°C with 660 mm of mean annual precipitation. FAB2 was designated to evaluate the influence of multiple dimensions of diversity and composition on long-term community processes and ecosystem function and resilience.

FAB2 is a tree diversity experiment established in 2016-2017 in an abandoned old field previously dominated by herbaceous species and fenced to exclude large mammalian herbivores. The experiment covers approximately 6.5 ha and has trees planted 1 m apart in 148 small plots (100 m^2^) and 30 large (400 m^2^) plots using twelve species with replicated monocultures and two-, four-, six-, and twelve-species polycultures plots (See more details in Cavender-Bares et al. in review) arrangement four experimental blocks. These species represent eight deciduous angiosperms (i.e., *Acer negundo*, *A. rubrum*, *Betula papyrifera*, *Quercus alba*, *Q. ellipsoidalis*, *Q. macrocarpa*, *Q. rubra*, *and Tilia americana*) and four evergreen gymnosperms (i.e., *Juniperus virginiana*, *Pinus banksiana*, *P. resinosa*, and *P. strobus*). For this study, we divide large plots into four small plots according to the tree locations within the plots to increase the number of replicates and standardize plot size and maximum tree density. The height and diameter of each tree in the experiment has been measured each year at the end of the growing season.

### Tree wood volume, productivity, and net biodiversity effects

Using the annual inventories of tree diameter and height that coincided with the study period (i.e., 2021 and 2022), we estimated wood volume for each tree, plot-level wood volume, annual tree and plot wood productivity, and the net biodiversity effects (NBE, *sensu* Loreau and Hector (2001), also referred to as overyielding) on annual wood productivity. Specifically, we estimated tree wood volume (m^3^) using the geometric formula for conoid and conoidoid volume and/or their joint sums. For trees less than 1.3 m, we used conoid volume (CV):

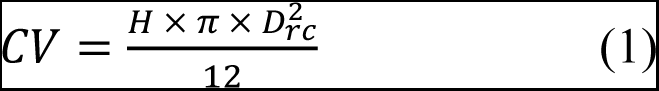

where *H* is the height of the tree from the root collar to the apical meristem, and *D* is stem diameter at the root collar (*rc*). For trees greater than 1.3 m, we used conoidoid plus conoid volume (CCV) as follows:

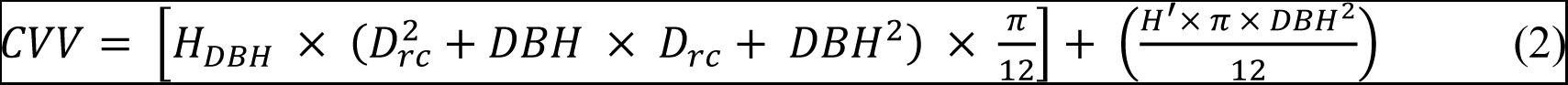

where *H_DBH_* is 1.3 m, *DBH* is diameter at breast height, and *H’* is the additional height above 1.3 m to the apical meristem. Annual tree wood productivity (AWP_tree_, m^3^ y^-1^) was then estimated as the difference in tree volume between 2022 and 2021 inventories divided by the number of days between inventories. Plot wood volume and annual wood productivity (AWP_plot_, m^3^ y^-1^) were estimated as the sum of tree volume and AWP_tree_ per plot, respectively. Using the AWP_tree_ estimations, we estimate NBE on productivity following Loreau and Hector (2001) as:

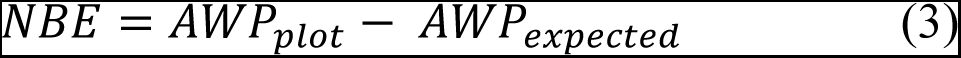

where AWP_expected_ is the expected annual plot productivity calculated as

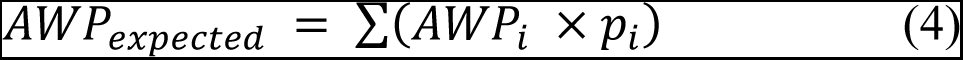

where AWP is the mean AWP_plot_ of the species *i* from monocultures that were planted before 2019, and *p* is the proportion of trees of the species *i* planted in the mixture. Our NBE estimate here includes tree mortality as part of the effect of diversity, but excludes trees replanted in 2022 to avoid potential increases in biomass due to the addition of trees. Trees at the border of the plots were excluded from these estimations to avoid the edge effect leading to an effective planting area of 64 m^2^.

### Diversity metrics

We estimate multiple dimensions of diversity associated with taxonomic (i.e., species richness), phylogenetic, and functional diversity though Hill numbers following Chao et al. (2014). Diversity metrics are all calculated such that they increase with numbers of species in the forest plot. Species richness was estimated as the effective number of species in the plot. Phylogenetic diversity (*PD*), was estimated using the effective total branch length using a pruned phylogenetic tree derived from Smith and Brown (2018). Finally, functional diversity (*FD*) was estimated as the effective total functional distance between species using functional traits related to growth, structure, and light strategies or requirements (i.e., wood density, leaf mass area, maximum tree height, relative growth rate, and shade tolerance) (Table S1). We computed these metrics of diversity using functions ‘*hill_taxa’*, ‘*hill_phylo’*, and ‘*hill_func’* of the hillR package (Li 2018) of R (R Core Team 2023) applying a Hill order *q =* 0. The phylogenetic tree was accessed through the ‘*phylo.maker’* function of the V.PhyloMaker package (Qian and Jin 2016) of R using the third scenario.

### Taxonomic, phylogenetic, and functional variability

Variability metrics were calculated to increase with taxonomic, phylogenetic, or functional divergence among species within a plot but to be independent of species richness. We used the phylogenetic species variability method proposed by Helmus et al. (2007), variability herein, to quantify the degree of dissimilarity within communities as a function of taxonomic, phylogenetic, or functional similarity. This metric, which is explicitly designed to be independent of species richness, is computed on phylogenetic trees or dendrograms, and provides a range of values between 0 (e.g., high relatedness or similarity among species) to 1 (e.g., low relatedness or similarity among species). The metric describes the degree of difference among members of a community, regardless of how many species are included. Similar to *PD* above, we used a pruned phylogenetic tree derived from Smith and Brown (2018) to estimate phylogenetic variability. We calculated functional variability using a dendrogram constructed from a multivariate Euclidean distance matrix of functional traits (Table S1) forced as a phylogeny.

Finally, we derived taxonomic variability by using a hierarchical clustering tree of taxonomic distinctness (Clarke and Warwick 1998) considering order, family, genus, and species. The clustering tree of taxonomic distinctness resembles the pruned phylogenetic tree (Figure S1), but provides a reasonable comparison that does not incorporate time-calibrated molecular branch length distances among taxa. We excluded monocultures from our analyses of variability given that the metric is undefined for a single species in the absence of measured intra-specific variability. This metric variability was estimated using the function ‘*psv*’ from *picante* package (Kembel et al. 2010), and the taxonomic distinctness using ‘*taxa2dist*’ from *vegan* package (Oksanen et al. 2022).

### LiDAR data collection and processing

We collected LiDAR data across the FAB experiment eight times during 2022 on the following days of the year: 100, 138, 163, 188, 215, 250, 261, and 297. Flights were conducted throughout the growing season spanning the entire favorable period for growth based on cumulative growing degree days (Figure S2), starting prior to leaf out and extending until after leaf senescence for the deciduous species. Data were collected using a Zenmuse L1 sensor onboard an Uncrewed Aerial Vehicle (UAV) DJI Matrice 300. This sensor integrates a Livox LiDAR module, an inertial measurement unit, and an RGB camera on a 3-axis stabilized gimbal. The LiDAR module has a conic footprint on ground with a field of view of 77.2° vertical and 70.4° horizontal, enabling it to capture multiple returns. All the surveys were conducted using autonomous flights programmed to take place at a speed of 8 m/s at 50 m above ground. These surveys were done in an area of ∼5.8 ha with 85% overlap between sidetracks allowing capture of dense point clouds (∼ 2100 point/m^2^). During these flights, we also collected static GNSS data using an Emlid Reach RS2+ receiver to enable kinematic corrections or post-processing corrections in case of signal loss. The data collected from the LiDAR sensor and GNSS receiver were processed in DJI Terra, delivering true color point clouds using the optimization option for manufacture strip alignment. The resulting point clouds were then processed through BayesStripAlign 2.24 to ensure a proper alignment among flight lines.

We treated the resulting point clouds in a series of steps to ensure data quality. First, points were classified as ground and non-ground and then normalized by height (Appendix 1). These procedures were done using ‘*lasthin’,* ‘*lasground’*, and ‘*lasheight’* of LAStools (Isenburg 2014). Then, we used a multi-polygon layer of the spatial limits of the experiment (e.g., Cavender-Bares et al. in review Figure 2) with a buffer of -0.5 m from the edge to segment point clouds. This buffer was applied to exclude trees planted at the edge similar to a previous section. We slightly aligned the multi-polygon layers within each survey using six long-term ground control points along the experiment to ensure spatial consistency between geospatial products.

**Figure 2.**
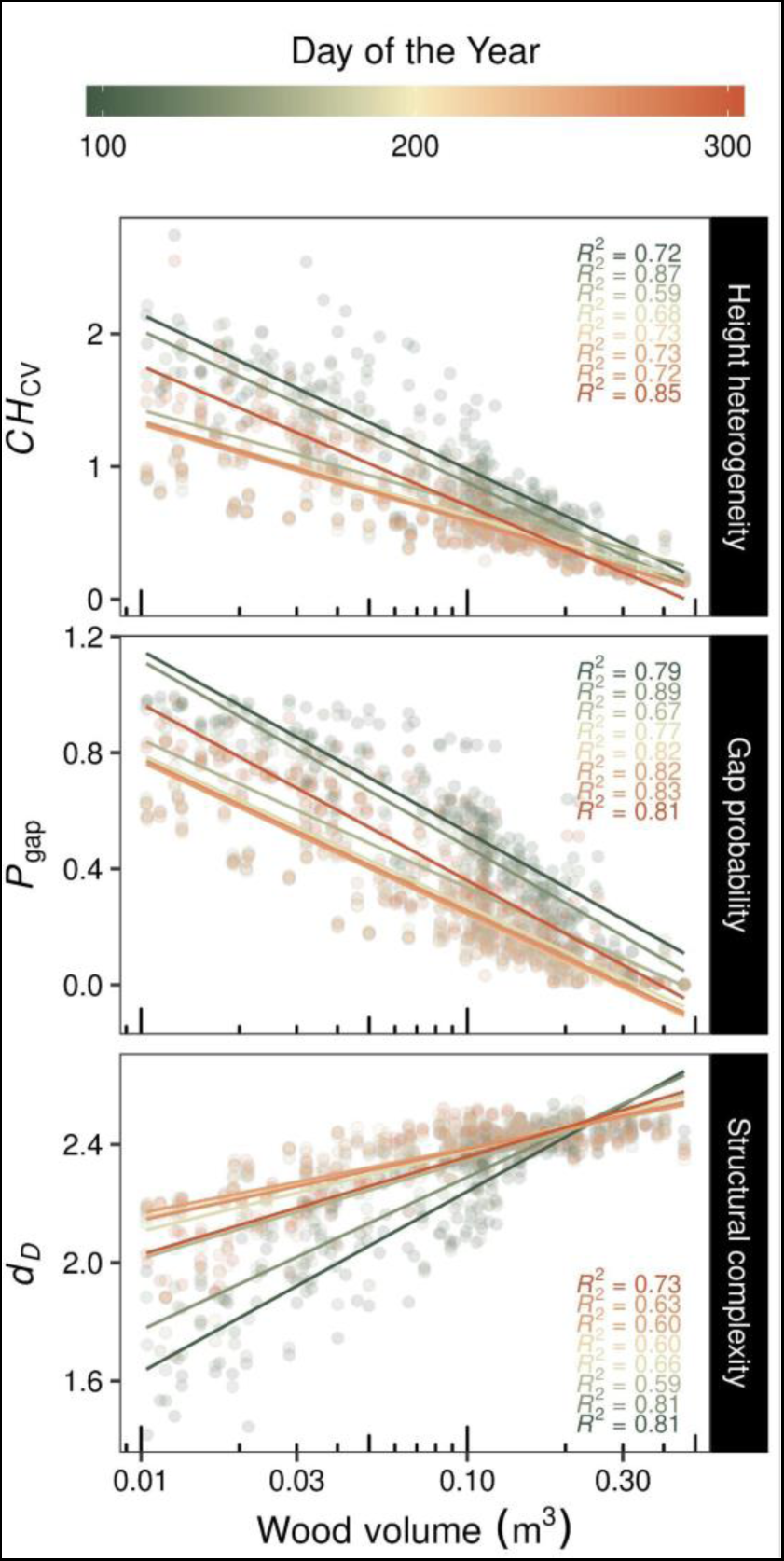
Influence of plot wood volume on LiDAR-derived metrics at different observation periods through the growing season. *CH*_CV_ describes the coefficient of variation of canopy height, *P*_gap_ the gap probability, and *d*_D_ the fractal dimension. Each point represents a plot within an area of 64m^2^. An extended figure with the significance of the regression models is presented in Figure S3.

The segmented point clouds were then filtered by noise using the Statistical Outliers Removal (SOR) algorithm considering 30 neighboring points and a threshold value equal to the 97.5% quantile. The filtering of noisy points was applied just on points above 0.25 m from the ground considered as vegetation. Once the segmented point clouds were filtered, we decimated them further to reduce the redundancy of points by selecting a point within a voxel grid of 1 cm of resolution. The filtering of points by noise and decimate procedure were done using the *lidR* package (Roussel et al. 2020).

### LiDAR-derived metrics

We derived three metrics to characterize the canopy heterogeneity, cover, and forest complexity from point clouds of each plot. Overall, we estimated: i) the coefficient of variation of canopy height (*CH*_cv_), ii) gap probability (*P*_gap_), and iii) the fractal dimension (*d*_D_), which integrate the canopy height heterogeneity, the light interception according to forest openness, and tridimensional arrangements of plant structures, respectively. We estimated these metrics only on plots with a canopy height higher than 1.5 m at the eight times that were surveyed, and on plots where the majority of trees were planted before 2019 (i.e., mean planted year per plot) (*n =* 170). Specifically, we computed *CH*_cv_ by estimating the coefficient of variation from cells of Digital Canopy Models from the plots with a 0.1 m of resolution. Likewise, we estimated *P*_gap_ by dividing the number of returns from the ground (i.e., < 0.25 m above the ground) by the total number of returns from the plot. Finally, we derived the *d*_D_ through the box-counting method as presented in Guzmán et al. (2020), but computed using Hill diversity (Jost 2006). For this, a point cloud from a plot can be covered using a voxel (*N*_1_ = 1) of edge size *S*_1_ (m), but as the voxel edge size (*S*) is reduced (*S*_1_ > *S*_2_ > *S_n_*), the number of voxels required (*N* > 1) to cover it increases. *N* increases as a power function, thus their slope *(d*) and intercept (*a*) can be computed using a linear model, as follows:

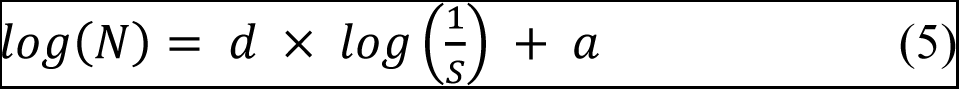

In order to incorporate Hill diversity into this formulation, *N* is replaced by estimating the Hill diversity (*D*) at each scale (i.e., *S*) following:

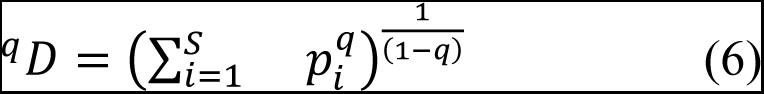

where *p*_i_ is the proportion of points that fall in a given voxel, and *q* are the different Hill orders (i.e., *q =* 0, *q =* 0.999, *q =* 2). This procedure allows us to compute different levels of fractal dimensions associated with the Hill orders (*d*_D_). In that regard, *d* resembles the box-counting dimension when *q* = 0, *d* resembles the information dimension when *q* approximates 1, and *d* resembles the correlation dimension when *q* = 2. Here, we focused our attention when *q* approximates 1 as it may help to reduce the differences in point density among plots (Liu et al. 2022). We solved the previous linear model by applying Standardized Major Axis (SMA) regressions using the ‘*sma*’ function of the smart package in R (Warton et al. 2012). To avoid the apparent fractality of point clouds due to occlusion effects, we used a sequence of voxel edge sizes (*S*) between 0.1 and 2.25 m of edge length. The estimations of *d*_D_ through voxelization were done with the help of the *lidR* (Roussel et al. 2020) and *rTLS* packages (Guzmán et al. 2021).

### Seasonal structural stability

We used the inverse of the coefficient of variation as a descriptor of temporal variation of forest structural metrics derived from LiDAR, and thus their structural stability (*SS*) across the season. This descriptor was estimated for each LiDAR metrics (i.e., *CH*_cv_, *P*_gap_, and *d*_D_) as follows:

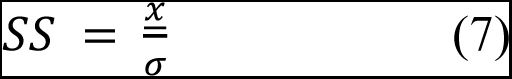

where 𝜎 is the standard deviation and *x* the mean of each metric from the eight surveys during the growing season. A higher value of *SS* describes a plot where its canopy height heterogeneity (*SS*_CHcv_), gap probability (*SS_Pgap_*), or structural complexity (*SS_d_*_D_) is stable across the growing season. We emphasize *SS_d_*_D_ in results as a comprehensive measure of structural stability given that it integrates the arrangements of forest structure in three-dimensions in contrast with *SS*_CHcv_ or *SS_Pgap_*.

### Data analysis

First, we performed separate linear mixed-models to evaluate the extent to which plot wood volume, diversity, and variability explain changes in LiDAR-derived metrics. These linear mixed models also consider the observation period effect and its interaction with the former parameters. For these models, we consider plots nested within experimental blocks as a repeated random effect. Second, we performed linear regressions between LiDAR-derived metrics at different observation periods and plot volume, diversity, and variability to determine how much of the variance (coefficient of determination, *R*^2^) is explained by changes in structure, diversity, and composition. Third, we applied linear mixed models to test how plot wood volume, diversity, and variability affect the structural stability of metrics of forest structure derived from LiDAR. Fourth, we assessed the influence of structural stability on NBE using linear regressions models as well. Finally, we performed structural equations models (SEMs) to assess pathways of how both diversity and variability affect forest structural stability, and how the latter in turn influences NBE (Figure 1d). These SEMs were evaluated using multiple dimensions of diversity and variability (i.e., taxonomic, phylogenetic, and functional) and were fitted by linear mixed-models using blocks as a random effect through the *piecewiseSEM* package (Lefcheck 2016). Linear mixed-models were fitted using the *lme4* package (Bates et al. 2015), while the linear regressions were performed using the *ggpmisc* package (Aphalo 2023) through ggplot2 (Wickham 2016).

## Results

### Relationships between wood volume and structural metrics derived from LiDAR

The relationships between plot wood volume and LiDAR-derived metrics of forest structure across the growing season reveal that increases in wood volume were strongly associated with reductions in canopy height heterogeneity (*CH*_CV_) and gap probability (*P*_gap_) and increases in structural complexity (*d*_D_) (Figure 2 and extended Figure S3). Variations of these trends among observation periods appear to emerge as a result of seasonal changes, where communities showed reduced heterogeneity and openness and increases in structural complexity close to the peak of the growing season (DOY 215 or August 15^th^) (Figure S4). The importance of seasonal changes in forest structure was also present in the linear mixed models (Table S2), but models also revealed an interaction between the observation period (i.e., DOY) and wood volume.

Moreover, our measures of forest structure stability across the growing season showed that increases in wood volume drive increases in structural stability of structural complexity (*SS_dD_*) and reductions in stability of gap probability (*SS_P_*_gap_), but do not influence structural stability of canopy height heterogeneity (*SS_CH_*_cv_) (Figure 3). Moreover, it appears that communities composed by gymnosperms and mixtures of angiosperms and gymnosperms present higher *SS_CH_*_cv_ and *SS_dD_* than communities composed solely of angiosperms (Figure S5). Structural stability in forest cover (i.e., *SS_1 - P_*_gap_) does not appear to be affected by angiosperm-gymnosperm composition.

**Figure 3.**
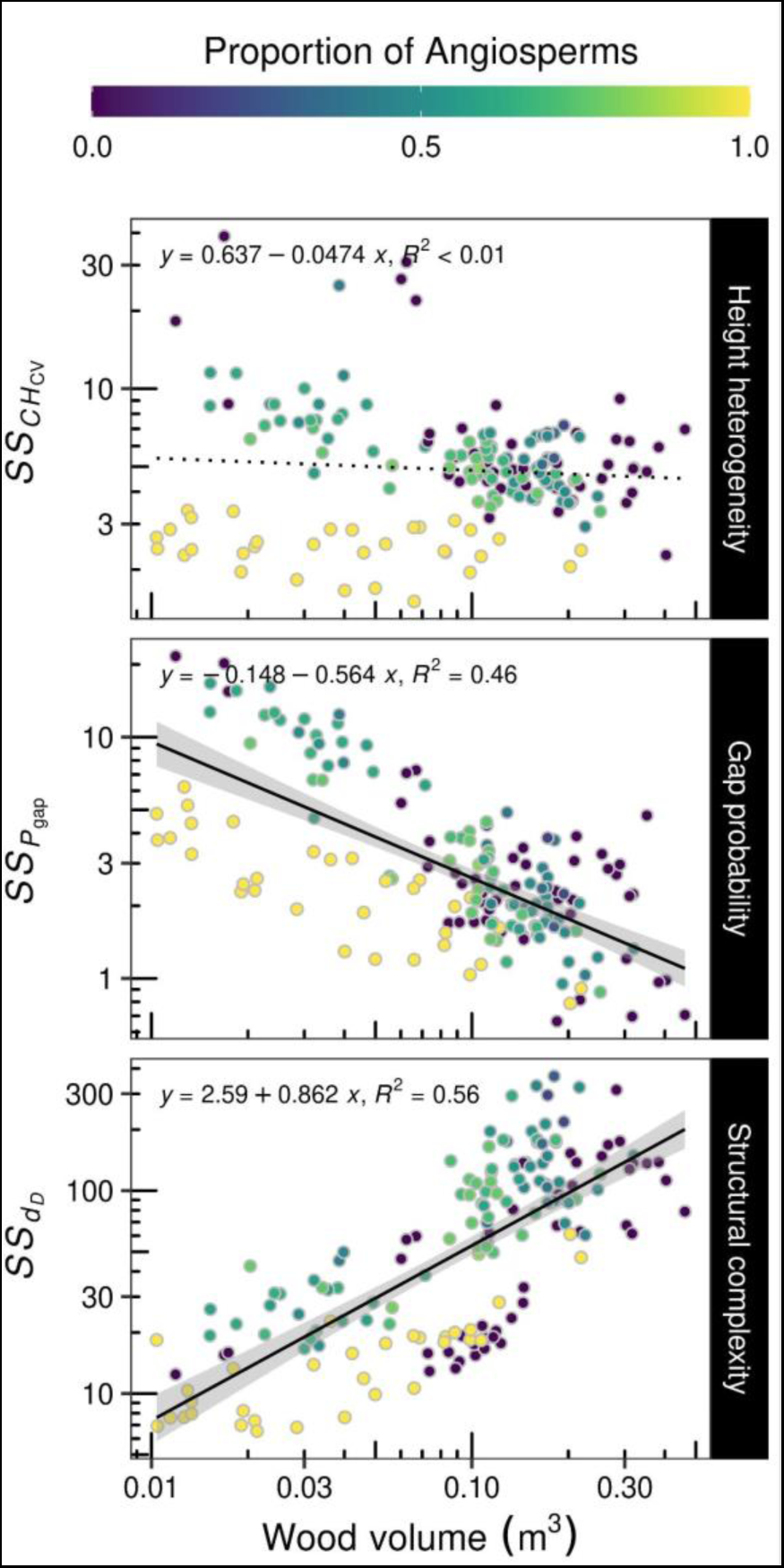
Effect of plot wood volume on the structural stability (*SS*) of LiDAR-derived metrics through the growing season. *CH*_CV_ describes the coefficient of variation of canopy height, *P*_gap_ the gap probability, and *d*_D_ the fractal dimension. Colors represent the proportion of angiosperms trees planted in each plot. Each point represents a plot within an area of 64m^2^.

*Effect of multiple dimensions of diversity on structural metrics derived from LiDAR* Analyses of the relationships between multiple dimensions of diversity and LiDAR-derived metrics revealed that increases in species richness, phylogenetic diversity (*PD*, calculated as effective total branch length), and functional diversity (*FD*, calculated as the effective total functional distance between species) tended to be associated with increases in canopy height *CH*_CV_ and *P*_gap_ (Figure 4, extended Figure S6, and Table S3). In most cases, the variance explained by these relationships is low (i.e., ≤ 30%) but appears to be better explained in observation periods close to the peak of the growing season. This result is also supported by the linear mixed models (Table S3), which indicate that observation periods have a significant influence on these LiDAR-derived metrics. The coefficient of determination of the regression models indicates that canopy height heterogeneity was the LiDAR-derived metric most tightly coupled with changes in diversity among plots. Moreover, the coefficient of determination also revealed that in most cases, changes in *FD* appeared to better explain changes in LiDAR-derived metrics than species richness or *PD*. An exception to this occurred when comparing the effects of richness and *FD* to *CH*_CV_, which were similar. When examining the seasonal structural stability, our results reveal that increases in diversity influence increases in the stability of forest structural metrics (with the exception of species richness and *FD* against *SS_dD_*) (Figure 5).

**Figure 4.**
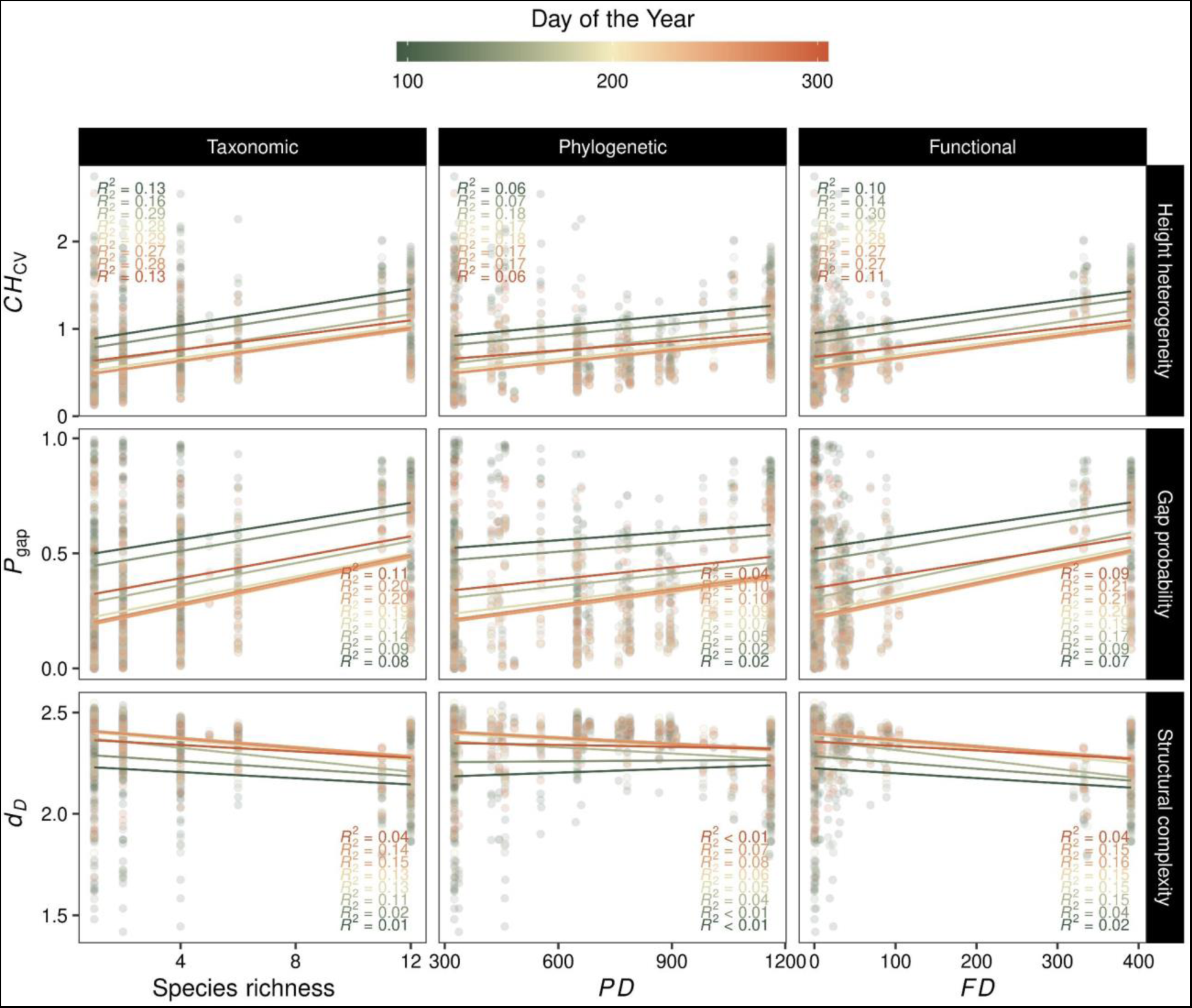
Influence of multiple dimensions of diversity on LiDAR-derived metrics at different observation periods during the growing season. *CH*_CV_ describes the coefficient of variation of canopy height, *P*_gap_ the gap probability, and *d*_D_ the fractal dimension. An extended figure with the significance of the regression models is presented in Figure S6.

**Figure 5.**
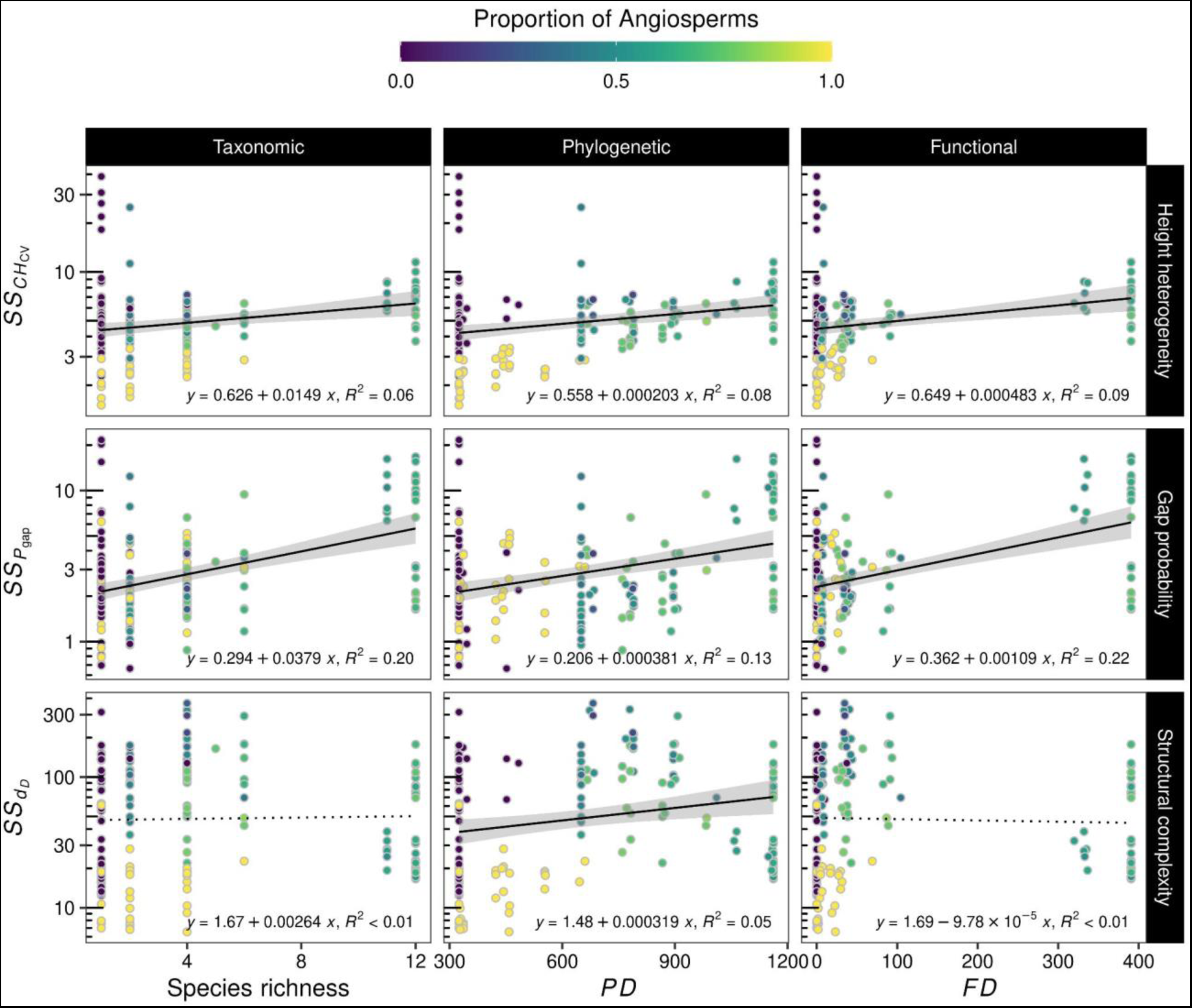
Effect of multiple dimensions of diversity on the structural stability (*SS*) of LiDAR-derived metrics through the growing season. *CH*_CV_ describes the coefficient of variation of canopy height, *P*_gap_ the gap probability, and *d*_D_ the fractal dimension. Most of these relationships were not statistically significant. Colors represent the proportion of angiosperms trees that were planted for each plot.

### Effect of multiple dimensions of variability on structural metrics derived from LiDAR

We found strong relationships between multiple dimensions of forest variability and structural metrics derived from LiDAR, revealing that communities composed of taxonomically, phylogenetically, and functionally dissimilar species had lower *CH*_CV_ and *P*_gap_ but higher forest *d*_D_ than plots composed of similar species (Figure 6 and extended Figure S7). In contrast to diversity where the relationships were better explained at the peak of the growing season (e.g., Figure 4), relationships of variability with forest structural metrics appear to be better explained in observation periods close to the beginning and end of the growing season. The variance explained by these relationships is also generally low ( ≤ 38%) and non-significant in most of the cases. These findings are supported by linear mixed models that revealed that observation periods and their interaction with multiple dimensions of variability have a significant effect on the evaluated forest structural metrics (Table S4). These relationships also indicate that functional variability within communities appears to better explain changes in structural metrics than phylogenetic or taxonomic variability (Figure 6 and Table S4). Interestingly, variability inversely influences forest structural metrics in comparison with diversity (e.g., Figure 4 vrs Figure 6) not just in the variance explained by linear models but also in the direction of trends. Similar to diversity (e.g., Figure 5), we found that variability influences structural stability such that increases in variability drive increases in the stability of canopy heterogeneity (*CH*_CV_) and forest complexity (*d*_D_), but not *P*_gap_ (Figure 7).

**Figure 6.**
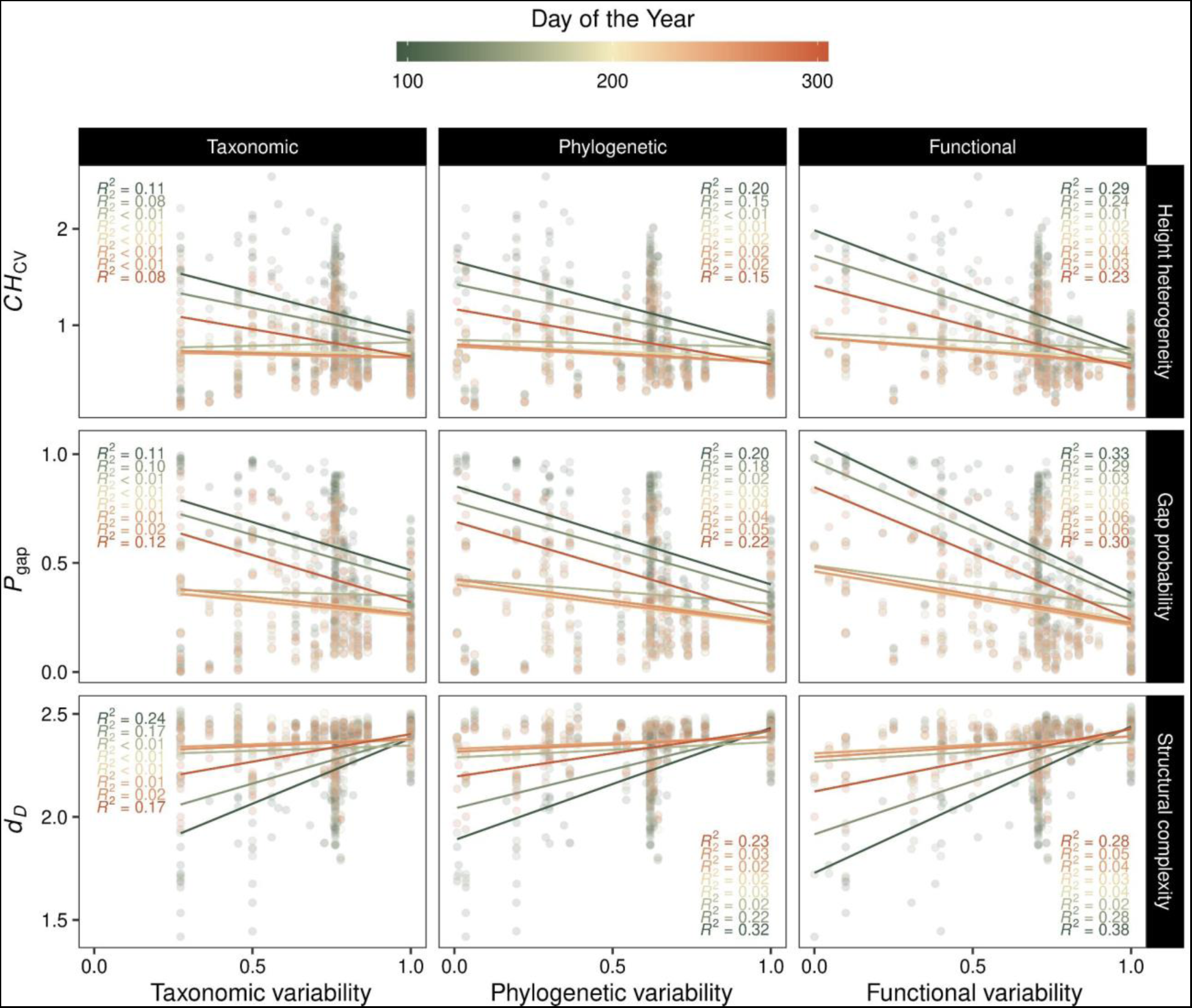
Influence of multiple dimensions of variability on LiDAR-derived metrics at different observation periods during the growing season. *CH*_CV_ describes the coefficient of variation of canopy height, *P*_gap_ the gap probability, and *d*_D_ the fractal dimension. An extended figure with the significance of the regression models is presented in Figure S6.

**Figure 7.**
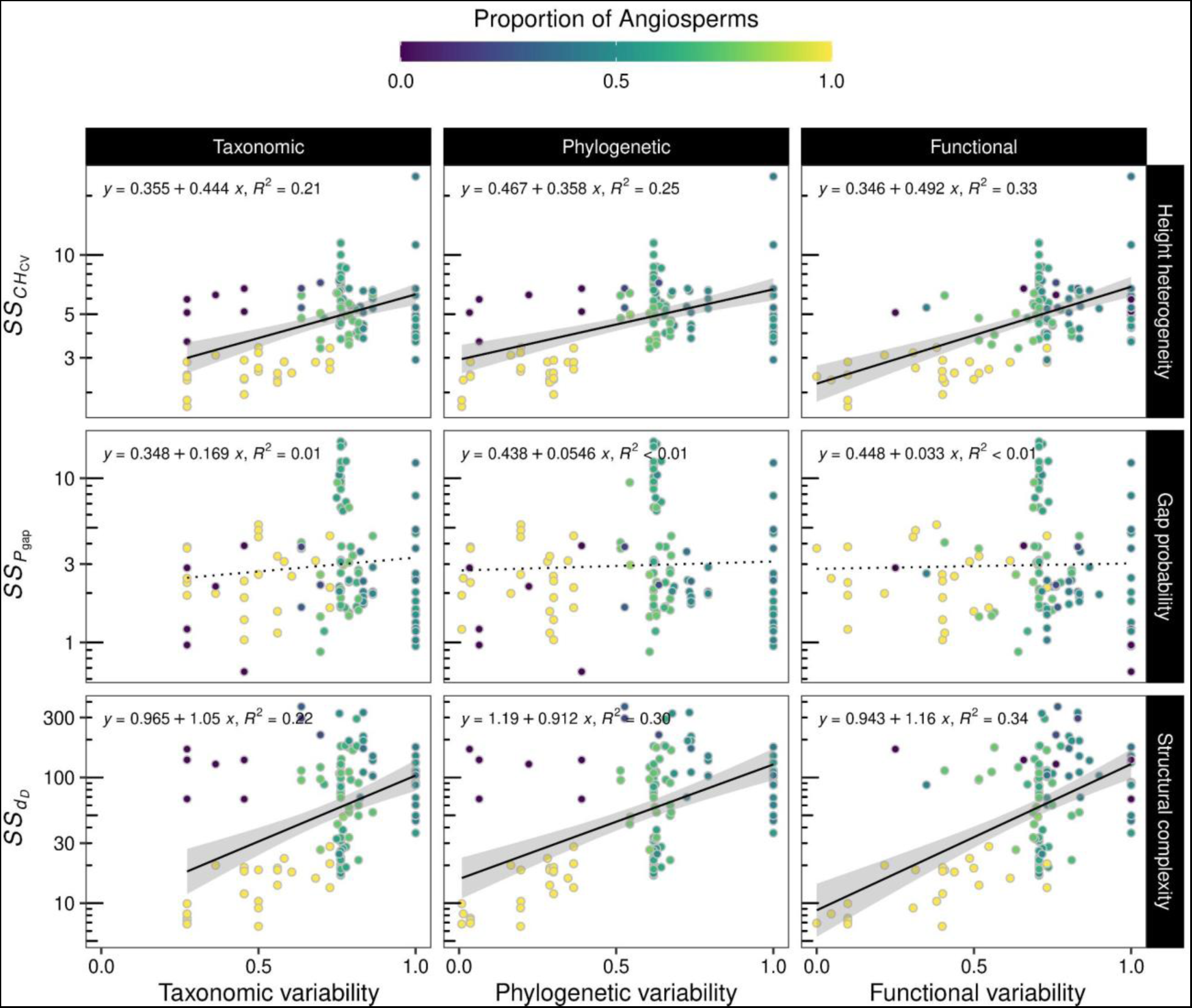
Effect of between different dimensions of variability on the structural stability (*SS*) of LiDAR-derived metrics through the growing season. *CH*_CV_ describes the coefficient of variation of the canopy height, *P*_gap_ the gap probability, and *d*_D_ the fractal dimension. Colors represent the proportion of angiosperms trees that were planted for each plot.

### Relationships with net biodiversity effect

We found significant relationships between the structural stability of LiDAR-derived metrics and net biodiversity effects (NBE). Structural stability of forest complexity (*SS_d_*_D_) and plant cover (*SS*_1-*Pgap*_), but not canopy height heterogeneity (*SS_CHcv_*) positively influence NBE (Figure 8). Increases in NBE were predominantly caused by complementarity effects rather than selection effects (Figure S8) according to the coefficient of determination. Moreover, NBE appeared to be negatively influenced by species richness and *FD* (Figure S9), but positively influenced by multiple dimensions of variability (Figure S10). In addition, plots composed by a higher proportion of angiosperms present lower NBE than those composed of a higher proportion of gymnosperms (Figure S11). Although diversity, variability, and the proportion of angiosperms trees planted have an influence on NBE, the variance explained by these relationships is not as high as the variance explained by the structural stability of forest complexity and gap probability (i.e., *SS_d_*_D_ and *SS_Pgap_*) (i.e., Figure 8).

**Figure 8.**
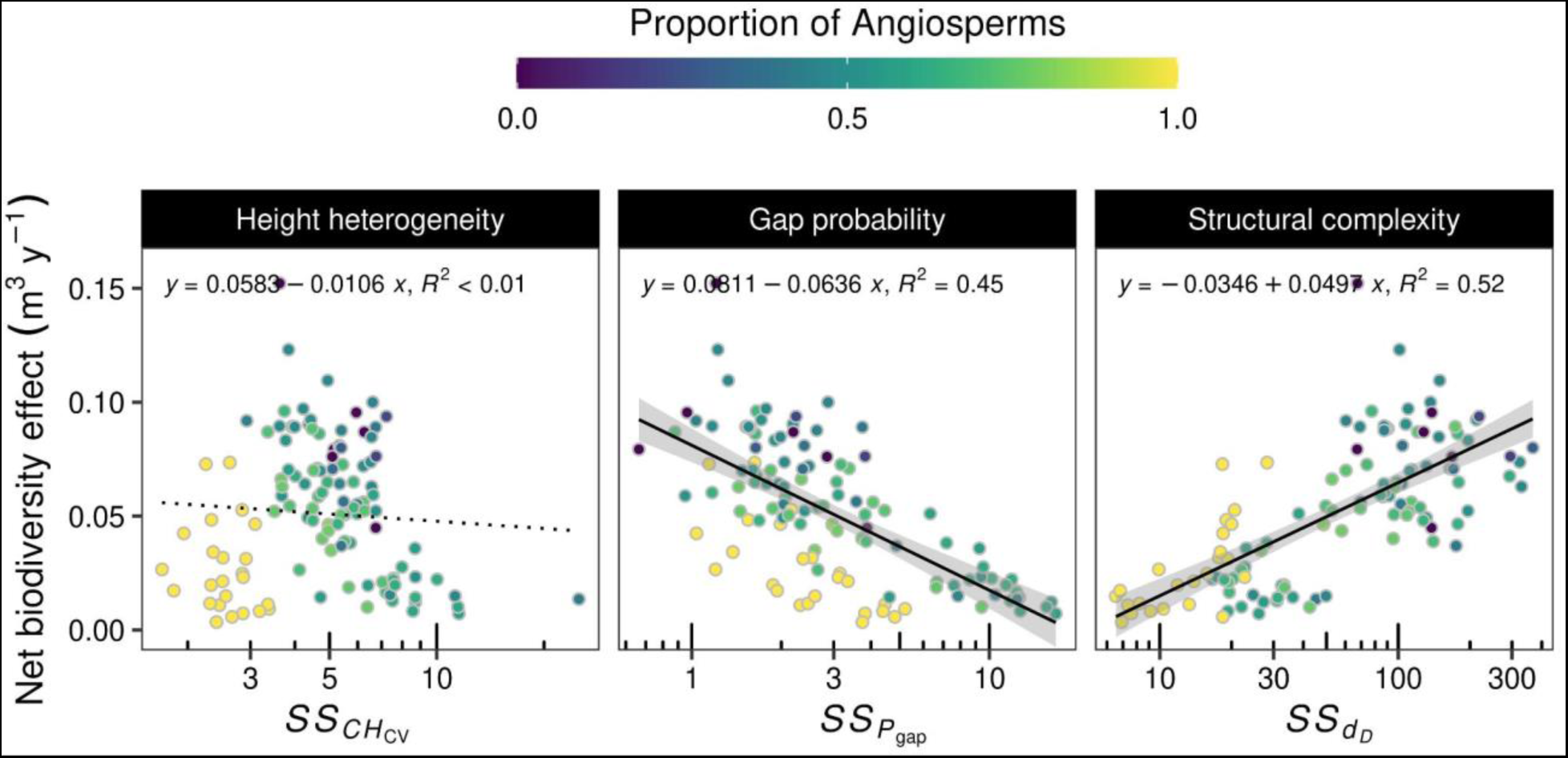
Influence of seasonal structural stability (*SS*) of LiDAR-derived metrics on the net biodiversity effect of annual wood productivity. *CH*_CV_ describes the coefficient of variation of the canopy height, *P*_gap_ the gap probability, and *d*_D_ the fractal dimension. Colors represent the proportion of angiosperms trees that were planted for each plot. An extended figure describing the complementary and selection effects is presented in Figure S8.

By integrating NBE in our structural equation models (SEMs), our results reveal that increases in NBE are mostly driven by increases in structural stability (Figure 9, and extended Figure S9). Both structural stability in forest complexity and gap probability (i.e., *SS_d_*_D_ and *SS_Pgap_*) are tightly coupled with NBE and appear to be more important in explaining variance in NBE than diversity or variability. These models based on *SS_d_*_D_ and *SS_Pgap_* tend to explain more than 61% of the variance in NBE (conditional *R*^2^). SEMs based on structural stability of canopy height heterogeneity (*SS*_CHcv_) do not explain more than 37% of the NBE variance when looking at multiple dimensions of diversity and variability. Diversity and variability, on the other hand, do not appear to have a consistent effect on NBE when compared among SEMs. Despite this, multiple dimensions of diversity and variability have a positive influence on metrics of structural stability, explaining up to 37% of their variance (range: 23% to 37%). The positive influence of diversity or variability shifts depending on the structural stability metric, being diversity more important to explain the *SS_Pgap,_* while variability to explain the *SS_d_*_D_. Diversity and variability tend to explain equally the variance in *SS*_CHcv_. Although slight differences appear when comparing between multiple dimensions of diversity and variability, structural stability metrics are better explained by phylogenetic diversity and variability, while NBE does not seem to be affected by the dimension of diversity and variability.

**Figure 9.**
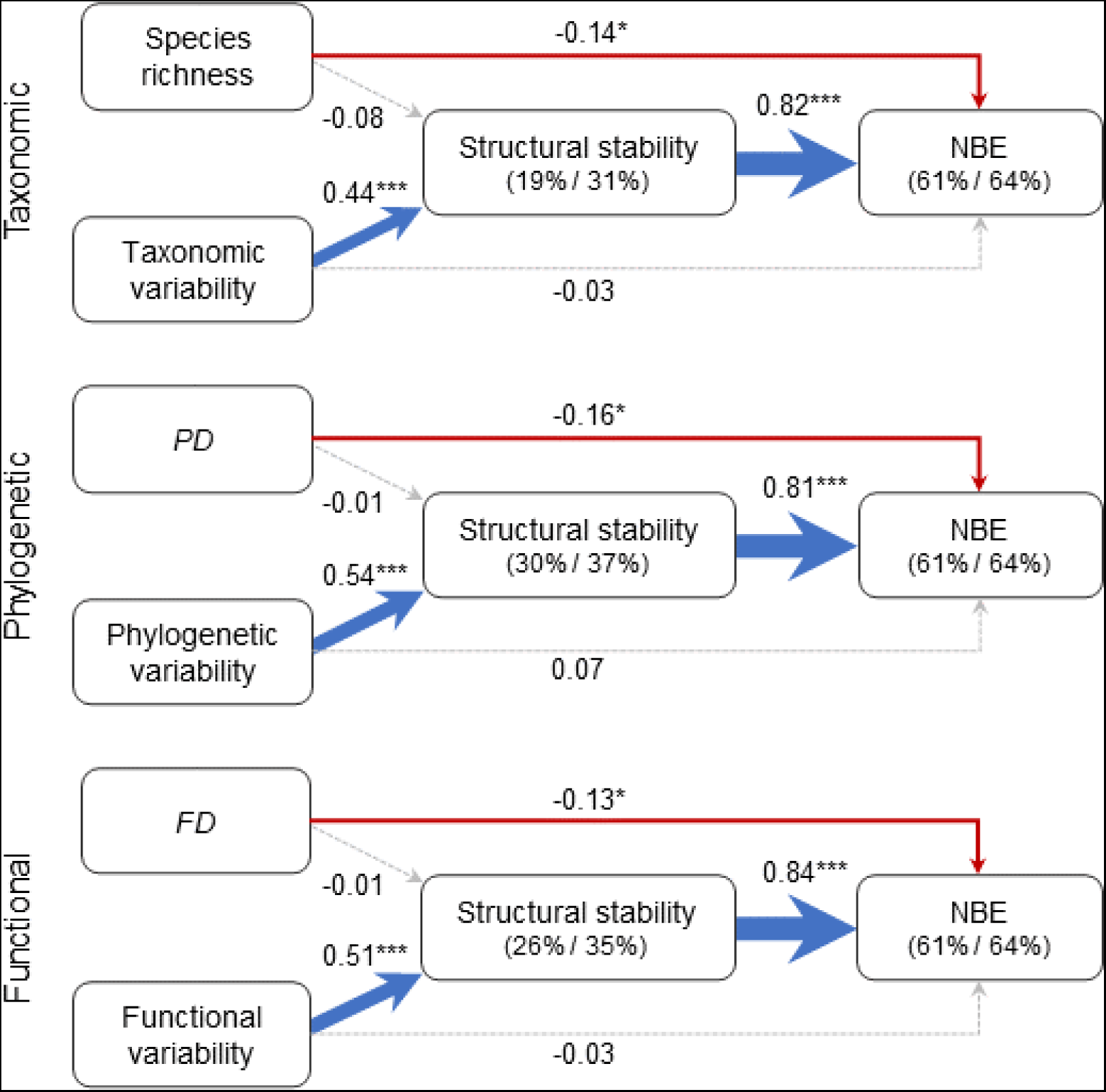
Structural equation models describing the paths of how multiple dimensions of diversity and variability influence the stability of forest structural complexity (i.e., fractal geometry) across the growing season, and in turn, affects the net biodiversity effect (NBE) of annual wood productivity (i.e., overyielding). Values next to the arrows represent the standardized coefficients and their significance, while values between parentheses coefficient of determination of marginal and conditional effects. The thickness of the arrows are defined by the standardized coefficients, while its color is positive (blue), negative (red), or non-effect (grey). An extended figure related to the other two metrics of forest structure (canopy height heterogeneity and gap probability) can be found in Figure S12.

## Discussion

In young stands of a large-scale tree diversity experiment, we show how multiple dimensions of forest diversity and variability influence structural metrics derived from LiDAR and their seasonal stability, which in turn, affect net biodiversity effects on productivity. This study is novel in investigating how multiple dimensions of diversity and variability (e.g., species dissimilarity) can drive the structural stability of forest communities through the growing season, highlighting emerging opportunities to understand the role of biodiversity on forest structure and its seasonality using remote sensing. Our work explains how forest structure, diversity, and variability influence LiDAR-derived metrics and highlights structural stability as an integrative metric that encompasses several drivers influencing net biodiversity effects on productivity in diverse forest communities.

### Relationships between forest volume on structural metrics derived from LiDAR

Our analyses revealed that variations in LiDAR-derived metrics are tightly coupled with changes in wood volume. These results indicate that LiDAR-derived metrics are reliable descriptors of current forest structure, and thus, the experimental diversity effects that can be derived from it. Nevertheless, our linear mixed models revealed that the interaction of observation period with structural variables can affect these relationships. Consequently, seasonal changes in forest communities are likely to influence our ability to predict remotely sensed structure. The interaction may result from differences in plant strategies (i.e., angiosperms - gymnosperms species) or timing of phenological events among species across the experiment, as well as variations in environmental conditions (e.g., wind) between the surveys that may affect our LiDAR estimations. Regardless of the influence of wood volume on the trajectories of LiDAR-derived metrics, our results show that increases in wood volume led to increases in the seasonal stability of structural complexity (i.e, *SS_d_*_D_) and plant cover (e.g., *SS*_1 *-Pgap*_). These outcomes might provide evidence that high seasonal structural stability appears as forests mature.

### Influence of multiple dimensions of diversity and composition on structural metrics derived from LiDAR and its seasonal stability

Our results support, to some extent, the premise that multiple dimensions of diversity influences LiDAR-derived metrics of forest structure. At the same time, our results challenge this premise because the direction of influence changes with taxonomic, phylogenetic, and functional similarity. Overall, the observed association of height heterogeneity (*CH*_CV_) with species richness is consistent with the canopy heterogeneity hypothesis (Torresani et al. 2020) in which tree species richness is expected to enhance canopy heterogeneity. We expand on this hypothesis with evidence that phylogenetic and functional diversity are also likely to enhance canopy heterogeneity. Moreover, increases in diversity also appeared to lead to communities that are more open and less structurally complex. This is an unexpected result because increases in diversity are expected to minimize space between trees through crown complementary while enhancing the light interception (Williams et al. 2017, 2021). However, in our study, canopy height heterogeneity and gap probability decay, and forest complexity increase when variability increases. These contradictory trends between diversity and variability observed from multiple dimensions (i.e., taxonomic, phylogenetic, and functional) might call into question the usefulness of the evaluated structural metrics derived from LiDAR to directly infer about forest biodiversity using single observation periods.

While diversity and variability show contrasting influences on LiDAR-derived metrics, we found that increases in both measures of biodiversity consistently showed positive influences on the stability of remotely sensed structural metrics across the season. Given that we did not find direct evidence that observed wood volume is associated with diversity (Figure S13) and as both measurements of biodiversity are not related to species abundance (e.g., species volume), our results provide tangible support that multiple dimensions of diversity and variability by themselves drive forest structural variation through the growing season, and thus its stability.

Forest diversity and variability can drive structural stability across the season due to the potential complementarity in structural strategies and the variability of phenological rhythms across the season among species integrated in forest mixtures. These factors may also explain why variability is more important in explaining differences in structural stability than diversity on forest complexity. Variability might be better to disentangle differences among species at low species richness levels (i.e., two species mixtures), which may be related to how contrasting their resource use strategies are, particularly over time. In this regard, communities composed of phylogenetically- or functionally-distant species result in higher structural stability than communities composed of similar species. In addition, our comparisons of multiple dimensions of diversity and variability indicate that the phylogenetic metrics (i.e., *PD* or phylogenetic variability) and, to some extent, functional metrics (i.e., *FD* or functional variability) are more important in describing changes in structural stability across the season than the taxonomic metrics (i.e., species richness or taxonomic variability) (e.g., Figure 9 or Figure S12).

### Remotely sensed seasonal structural stability as an integrative metric to describe net biodiversity effects

Our results indicate that seasonal structural stability, promoted by multiple dimensions of diversity and variability, is likely to enhance net biodiversity effects on productivity in forest communities. Niche complementarity in resource use among species is often the mechanism proposed for explaining overyielding (Kelty 1992, Tilman et al. 2001). Several mechanisms have been documented that explain enhanced net biodiversity effects on biodiversity, including tree crown complementary (Williams et al. 2017), increased light interception and efficiency of light use (Williams et al. 2021), or temporal resource partitioning (Lu et al. 2016). Our remote estimates of seasonal structural stability integrate all these potential complementarity mechanisms and help to explain net biodiversity effects. For instance, how well tree crowns complement each other, and thus enhance NBE, is indirectly described by structural complexity (*d*_D_), because assemblages that have higher and more stable complexity (Guzmán et al. 2020) are those with structures that homogeneously fill in their available three-dimensional space over time. Likewise, gap probability (*P*_gap_) and its stability capture enhanced light interception, given that *P*_gap_ represents the probability that laser pulses will be intercepted by trees before reaching the forest floor. We further expect temporal resource partitioning to be captured by LiDAR-based measures of seasonal structural stability, which integrate information about the phenological variation among species that is linked to intrinsic growth strategies (i.e., fast- or slow-growing). The importance of temporal resource partitioning for biodiversity effects on productivity is supported by Lu et al. (2016) who suggest that differences in leaf phenology and shade tolerance leads to increases in overyielding in evergreen-deciduous mixtures, but not deciduous-deciduous mixtures. Evergreen-deciduous mixtures (i.e., gymnosperms-angiosperms in our case) are expected to capture and use resources differently across the season, helping to explain why complementarity, and not selection effects, is more tightly coupled with structural stability (e.g., Figure S8). Distantly related groups of species, in general, tend to capture and use resources in contrasting ways, reducing competition and contributing to their complementarity. In long-term grassland experiments, for example, communities composed of distantly related species buffer ecosystems against environmental variation resulting in greater stability of productivity (Cadotte et al. 2012). Moreover, canopy height heterogeneity (*CH*_CV_) only represents the variation at the top of the canopy but does not describe the three-dimensionality of forest structure, which may explain why this metric is not as clearly linked to net biodiversity effects on productivity.

### Remote sensing of forest structural stability as tool to evaluate ecological process

Remotely-sensed forest structural stability through the season appears to integrate several drivers of overyielding. Therefore, seasonal structural stability may help to discern the role of diversity in other ecological processes within forest communities such as alteration of microclimates, forest resilience, and ecosystem functions that generate important services. For instance, given the strong influence of forest structure on microclimates (Potter et al. 2001), it is likely that structural stability promoted by diversity and variability alter temporal variation of microclimates within forest communities by buffering them from environmental extremes.

Temporal variation of microclimates product to structural stability is likely to drive changes in below-ground diversity (Lang et al. 2023) and therefore the availability of nutrients. In addition, structurally stable and complex communities may create optimal conditions that can be used by other plant species to colonize available spaces (Walter et al. 2021) as well as to enhance below-ground diversity (Lang et al. 2023). Currently, there is a large number of metrics that can be derived from LiDAR (e.g., Atkins et al. 2023a); many of which are affected by the spatial scale (Atkins et al. 2023b) or are redundant in terms of their ecological meaning. Therefore, the future integration of LiDAR with ecology should focus on metrics that are not dependent on spatial or temporal scales as well as on metrics that help to evaluate multiple dimensions of biodiversity.

## Conclusions

Here we provide empirical evidence that multiple dimensions of diversity and variability (e.g., species disimilarity) have an important influence on structural metrics derived from UAV-LiDAR. Most importantly, we show that increases in diversity and variability of forest communities have a positive influence on the temporal stability of forest structure throughout the growing season. We further demonstrate that high structural stability over the season of forest structural complexity (i.e., fractal geometry) and plant cover (i.e., 1 - gap probability) enhances net biodiversity effects of productivity due to species complementarity. Our findings highlight remotely sensed structural stability as an integrative metric that encompasses several drivers of net biodiversity effects of forest productivity. These findings demonstrate the potential couple remote sensing and ecology to study forest stability to unravel its role into ecological processes through three-dimensional information.

## Author Contributions

JAGQ and JC-B designed the research. JAGQ conducted the UAV-LiDAR surveys with the assistance of JC-B. JAGQ processed, analyzed, and visualized the data. MHP drafted the conceptual figure with inputs from JAGQ and JC-B. JAGQ drafted the manuscript with inputs from MHP, LJW, and JC-B. Funding sources come through JC-B. All authors contributed intellectually to the project and to editing-directions of the manuscript.

## Acknowledgments

We acknowledge that this study was conducted on Dakota and Ojibwe land. We thank Cathleen Lapadat, Allison Scott and Erica Halek, and for helping to establish the experiment and conducting annual forest inventories. We thank Sarah Hobbie and Rebecca Montgomery for contributions to the design and establishment of the experiment, and Matthew Kaproth for seed sourcing and seed collection. We thank the NSF LTER grant for Cedar Creek Ecosystem Science Reserve (DEB:1831944) and the NSF ASCEND Biology Integration Institute (DBI: 2021898) for supporting the work.

## Open research statement

Forest inventory data will be available at the Environmental Data Initiative and Cedar Creek Data Catalog upon publication. The processed point clouds from the UAV-LiDAR surveys are already in DRYAD, but will be released to the public upon publication. All the code used to process, analyze, and visualize data is openly available at GitHub (https://github.com/Cavender-Bares-Lab/UAV-LiDAR-Processing.git) and will be archived at Zenodo under version 1.0 upon publication.

