## Supplementary materials for "Seasonal structural stability promoted by forest diversity and composition explains overyielding"

**Appendix 1** - LiDAR ground classification and height normalization

We applied a series of steps to ensure an optimal classification of points as ground/non-ground and its posterior normalization by height. These steps were inspired, but not similar, to the procedure described by Mattie (2022) for the Zenmuse L1 LiDAR sensor which attempts to correct the 'thickness' of the point clouds using LAStools (Isenburg 2014). For this we first select low candidates for ground points using percentiles between 10% and 20% according to point height applying consecutive different grid sizes (20, 40, 60 cm). At each grid size these low candidate points were classified as model points (Class 8) using *lasthin*. On the resulting low candidates, we then run a ground classification through *lasground* in which low ground points (Class 2) are defined. This low ground classification procedure was defined using a 'coarse' and 'nature' settings avoiding spikes above 0.25 m for better performance. Using these low ground points we then run a height normalization through *lasheight* to replace point elevation by point height from the ground; providing height-normalized point clouds for further analysis.

**Table S1.** Characteristics and plant functional traits of species at the Forest and Biodiversity Experiment II. LMA describes the leaf mas area, and RGR the relative growth rate.

| Order | Family | Species | Wood density[1] | LMA[2] | Shade tolerance index[3] | Maximum height[4] (m) | RGR[5] |
|---|---|---|---|---|---|---|---|
| Sapindales | Sapindaceae | *Acer negundo* | 0.44 | 37.04 | 3.47 | 18.59 | 0.039 |
| Sapindales | Sapindaceae | *Acer rubrum* | 0.44 | 71.09 | 3.44 | 19.81 | 0.379 |
| Fagales | Betulaceae | *Betula papyrifera* | 0.48 | 77.88 | 1.54 | 11.58 | 1.272 |
| Pinales | Cupressaceae | *Juniperus virginiana* | 0.44 | 333.33 | 1.28 | 14.63 | 0.975 |
| Pinales | Pinaceae | *Pinus banksiana* | 0.4 | 243.9 | 1.36 | 22.25 | 2.288 |
| Pinales | Pinaceae | *Pinus resinosa* | 0.41 | 294.12 | 1.89 | 25.91 | 1.619 |
| Pinales | Pinaceae | *Pinus strobus* | 0.34 | 121.92 | 3.21 | 27.13 | 1.722 |
| Fagales | Fagaceae | *Quercus alba* | 0.6 | 81.21 | 2.85 | 24.38 | 0.735 |
| Fagales | Fagaceae | *Quercus ellipsoidalis* | 0.59 | 90 | 2.49 | 23.16 | 0.722 |
| Fagales | Fagaceae | *Quercus macrocarpa* | 0.58 | 92.74 | 2.71 | 21.64 | 0.547 |
| Fagales | Fagaceae | *Quercus rubra* | 0.56 | 84.2 | 2.75 | 24.99 | 0.505 |
| Malvales | Malvaceae | *Tilia americana* | 0.32 | 60.81 | 3.98 | 23.77 | 0.724 |

1

2

[3] Data from Niinemets and Valladares (2006)

[4] Maximum height (quantile 99%) of trees found in inventories between 2018 to 2022 from the Forest Inventory and Analysis National Program across Minnesota, US.

[5] Mean relative growth rate of species on monocultures between 2013 to 2016 from the first Forest and Biodiversity Experiment at Cedar Creek Ecosystem Science Reserve, Minnesota, USA (Kothari et al. 2021).

**Table S2.** Linear mixed model statistics for comparing the wood volume and day of the year (DOY), and their interaction on LiDAR derived metrics. Values of the fixed effects represent the *F-ratio* of Analysis of Variance using Type II Wald F-tests, values in parentheses are the degrees of freedom, and the asterisks next to each value the significance. $\sigma^2$ describes the variance of the model, $\tau_{00}$ the variance of the random effects, and ICC the interclass correlation coefficients.

| Predictors | LiDAR-derived metric | | |
|---|---|---|---|
| | Canopy height heterogeneity ($CH_{CV}$) | Gap probability ($P_{gap}$) | Structural complexity ($d_D$) |
| **Fixed effects** | | | |
| Volume | **602.41***** (1 / 168) | **1460.21***** (1 / 168) | **731.44***** (1 / 168) |
| DOY | **364.12***** (7 / 1176) | **288.37***** (7 / 1176) | **144.07***** (7 / 1176) |
| Interaction | **25.54***** (7 / 1176) | **10.28***** (7 / 1176) | **135.70***** (7 / 1176) |
| **Random effects** | | | |
| $\sigma^2$ | 0.02 | 0.01 | 0.00 |
| $\tau_{00}$ plot:block | 0.05 | 0.00 | 0.00 |
| ICC | 0.76 | 0.38 | 0.42 |
| Marginal $R^2$ | 0.77 | 0.83 | 0.75 |
| Conditional $R^2$ | 0.94 | 0.90 | 0.85 |

*p-value*: * <0.05; ** <0.01, *** <0.001

**Table S3.** Linear mixed model statistics for evaluating the effect of multiple dimensions of diversity, the day of the year (DOY), and their interaction on LiDAR derived metrics. Values of the fixed effects represent the *F-ratio* of Analysis of Variance using Type II Wald F-tests, values in parentheses are the degrees of freedom, and the asterisks next to each value the significance. ICC describes the intraclass correlation coefficients.

| Dimensions of diversity | Effects | Predictors | LiDAR-derived metric | | |
|---|---|---|---|---|---|
| | | | Canopy height heterogeneity ($CH_{CV}$) | Gap probability ($P_{gap}$) | Structural complexity ($d_D$) |
| Taxonomic | Fixed | Species richness | **41.92*** ** (1 / 168) | **20.74*** ** (1 / 168) | **7.03** ** (1 / 168) |
| | | DOY | **117.87*** ** (7 / 1176) | **274.62*** ** (7 / 1176) | **86.75*** ** (7 / 1176) |
| | | Interaction | **2.27* ** (7 / 1176) | 1.78 (7 / 1176) | 1.56 (7 / 1176) |
| | Random | ICC | 0.82 | 0.86 | 0.73 |
| | | Marginal $R^2$ | 0.27 | 0.24 | 0.14 |
| | | Conditional $R^2$ | 0.87 | 0.89 | 0.76 |
| Phylogenetic | Fixed | *PD* | **18.70*** ** (1 / 168) | **6.60* ** (1 / 168) | 0.62 (1 / 168) |
| | | DOY | **176.70*** ** (7 / 1176) | **276.82*** ** (7 / 1176) | **91.98*** ** (7 / 1176) |
| | | Interaction | 1.14 (7 / 1176) | **3.14** ** (7 / 1176) | **11.58*** ** (7 / 1176) |
| | Random | ICC | 0.84 | 0.87 | 0.75 |
| | | Marginal $R^2$ | 0.20 | 0.18 | 0.12 |

| | | | | | |
|---|---|---|---|---|---|
| | | Conditional $R^2$ | 0.87 | 0.89 | 0.78 |
| Functional | Fixed | *FD* | **32.42***** (1 / 168) | **14.78***** (1 / 168) | 3.86 (1 / 168) |
| | | DOY | **176.82***** (7 / 1176) | **275.21***** (7 / 1176) | **88.12***** (7 / 1176) |
| | | Interaction | 1.26 (7 / 1176) | **2.14*** (7 / 1176) | **4.05***** (7 / 1176) |
| | Random | ICC | 0.83 | 0.86 | 0.73 |
| | | Marginal $R^2$ | 0.24 | 0.22 | 0.13 |
| | | Conditional $R^2$ | 0.87 | 0.89 | 0.77 |

*p-value*: * <0.05; ** <0.01, *** <0.001

**Table S4.** Linear mixed model statistics for evaluating the effect of multiple dimensions of species variability, the day of the year (DOY), and their interaction on LiDAR derived metrics. Values of the fixed effects represent the *F-ratio* of Analysis of Variance using Type II Wald F-tests, values in parentheses are the degrees of freedom, and the asterisks next to each value the significance. ICC describes the intraclass correlation coefficients.

| Dimensions of variability | Effects | Predictors | LiDAR-derived metric | | |
|---|---|---|---|---|---|
| | | | Canopy height heterogeneity ($CH_{CV}$) | Gap probability ($P_{gap}$) | Structural complexity ($d_D$) |
| Taxonomic | Fixed | Species richness | 2.78 (1 / 115) | **2.61*** (1 / 115) | **10.67**** (1 / 115) |
| | | DOY | **251.65***** (7 / 805) | **330.15***** (7 / 805) | **80.75***** (7 / 805) |
| | | Interaction | **38.46*** (7 / 805) | **26.93***** (7 / 805) | **49.12***** (7 / 805) |
| | Random | ICC | 0.90 | 0.92 | 0.78 |
| | | Marginal $R^2$ | 0.19 | 0.21 | 0.22 |
| | | Conditional $R^2$ | 0.92 | 0.93 | 0.83 |
| Phylogenetic | Fixed | *PD* | **8.45**** (1 / 115) | **11.63***** (1 / 115) | **16.83***** (1 / 115) |
| | | DOY | **285.59***** (7 / 805) | **352.69***** (7 / 805) | **88.83***** (7 / 805) |
| | | Interaction | **59.15***** (7 / 805) | **36.62***** (7 / 805) | **65.54***** (7 / 805) |
| | Random | ICC | 0.91 | 0.92 | 0.79 |
| | | Marginal $R^2$ | 0.24 | 0.25 | 0.26 |

| | | | | | |
|---|---|---|---|---|---|
| | | Conditional $R^2$ | 0.93 | 0.94 | 0.85 |
| Functional | Fixed | *FD* | **14.23**** (1 / 115) | **18.03***** (1 / 115) | **21.87***** (1 / 115) |
| | | DOY | **353.78***** (7 / 805) | **447.28***** (7 / 805) | **100.06***** (7 / 805) |
| | | Interaction | **100.74***** (7 / 805) | **77.28***** (7 / 805) | **88.37***** (7 / 805) |
| | Random | ICC | 0.92 | 0.93 | 0.81 |
| | | Marginal $R^2$ | 0.28 | 0.29 | 0.30 |
| | | Conditional $R^2$ | 0.94 | 0.95 | 0.86 |

*p-value*: * <0.05; ** <0.01, *** <0.001

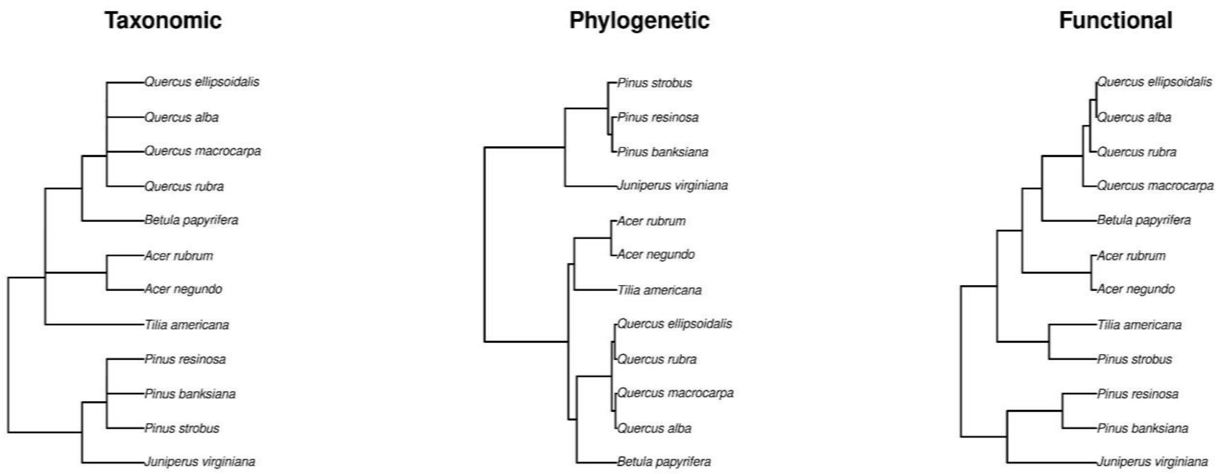


**Figure S1.** Taxonomic, phylogenetic, and functional trees of the twelve species present at the Forest and Biodiversity 2 tree experiment.

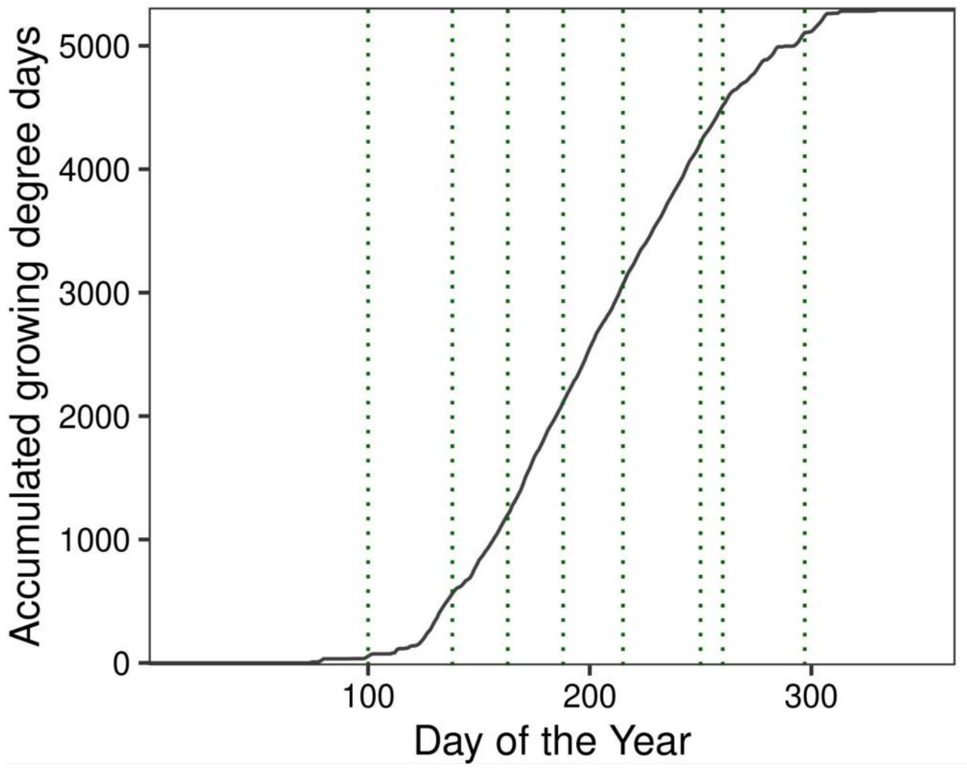


**Figure S2.** Accumulated growing degree days (base 15°C) for 2022 at the Cedar Creek Ecosystem Science Reserve (CCESR), Minnesota, USA. The estimated growing degree days were computed using meteorological measurements available from Seeley (2023).

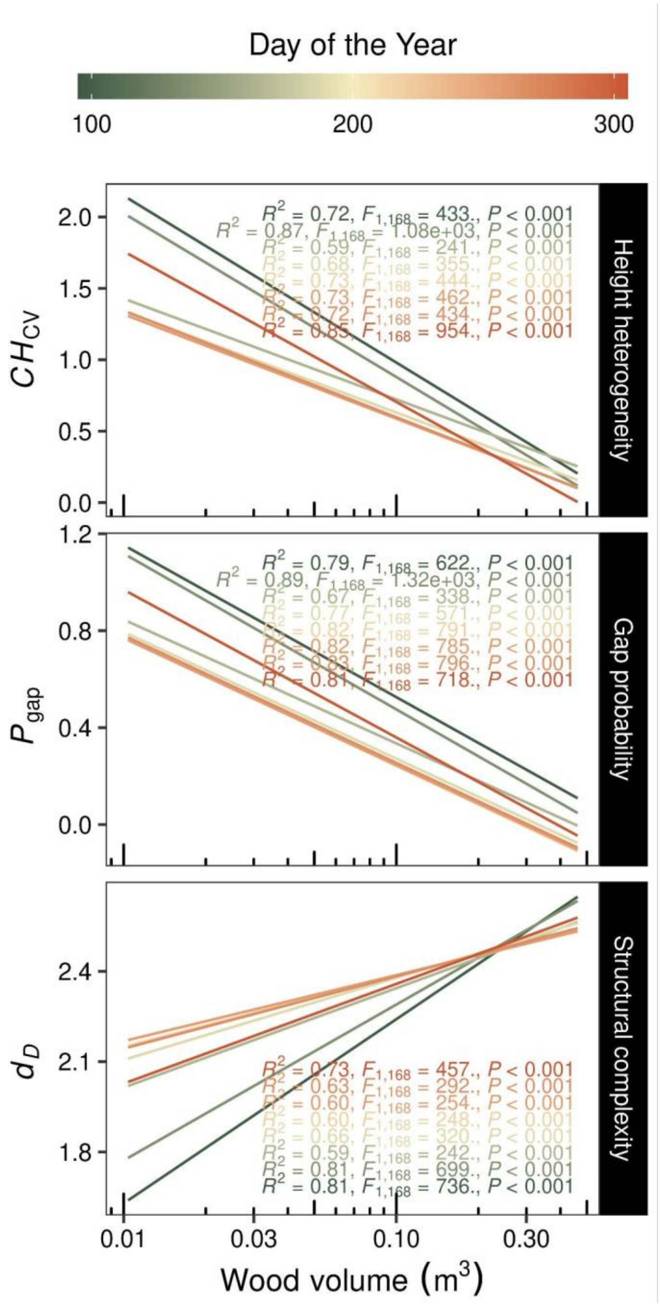


**Figure S3.** Influence of plot wood volume on LiDAR-derived metrics at different observation periods through the growing season. $CH_{\mathrm{CV}}$ describes the coefficient of variation of canopy height, $P_{\mathrm{gap}}$ the gap probability, and $d_{\mathrm{D}}$ the fractal dimension. The plot wood volume was derived from experimental plots within an area of $64\mathrm{m}^2$.

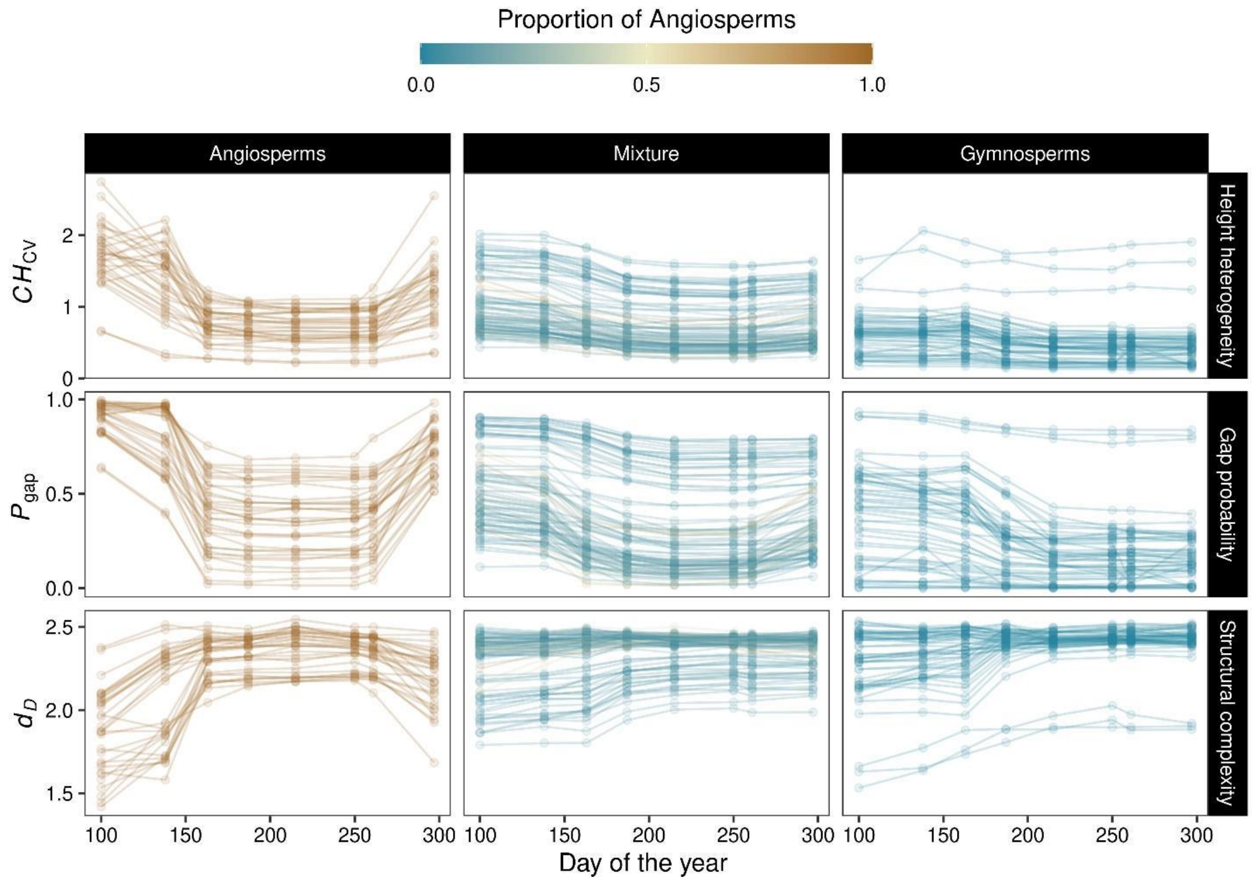


**Figure S4.** Temporal variation of the coefficient of variation of canopy height ($CH_{CV}$), gap probability ($P_{gap}$), and fractal geometry ($d_D$) of plots composed by angiosperms, gymnosperms, and mixture (i.e., angiosperms and gymnosperms) of species. Each line represents a forest plot, and each point an observation period.

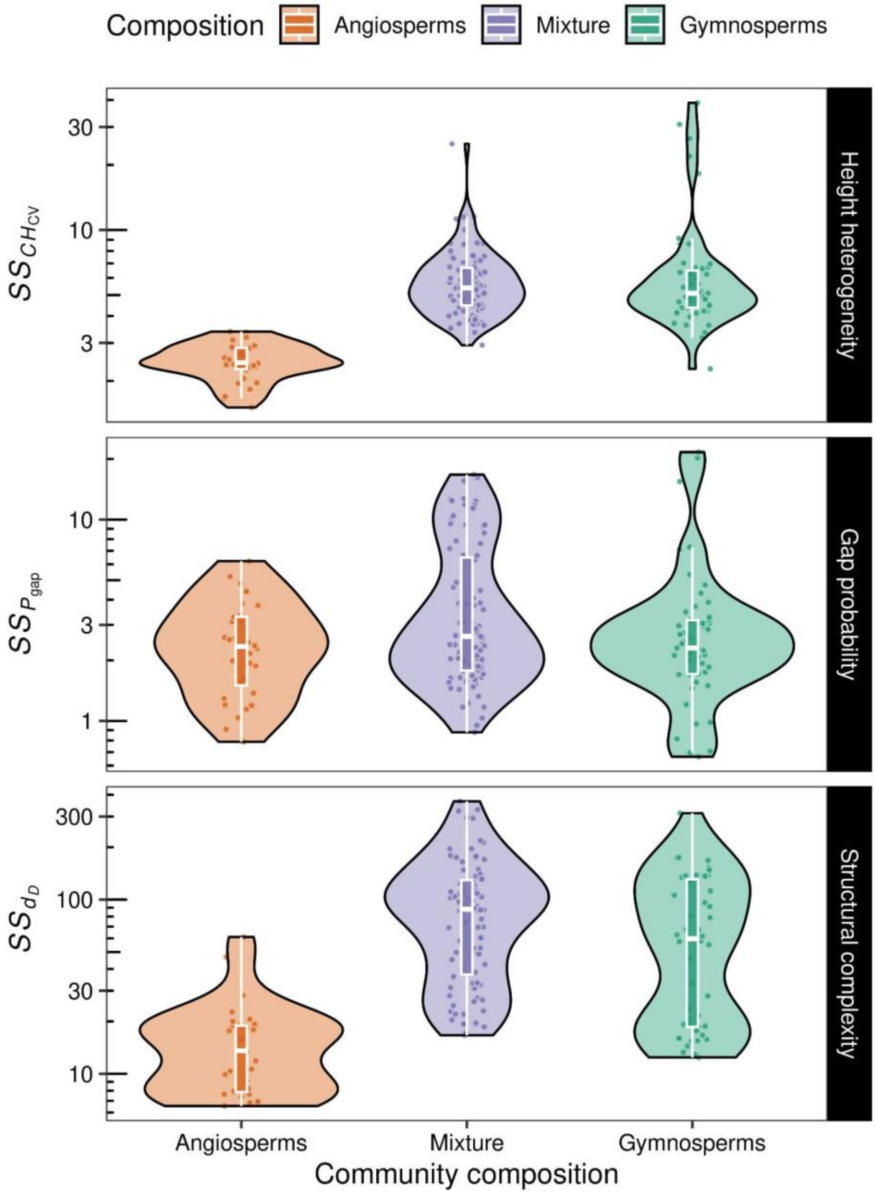


**Figure S5.** Comparison of seasonal structural stability (*SS*) of LiDAR-derived metrics among plots composed of angiosperms, gymnosperms, or mixtures (angiosperms and gymnosperms). $CH_{CV}$ describes the coefficient of variation of canopy height, $P_{gap}$ the gap probability, and $d_D$ the fractal dimension.

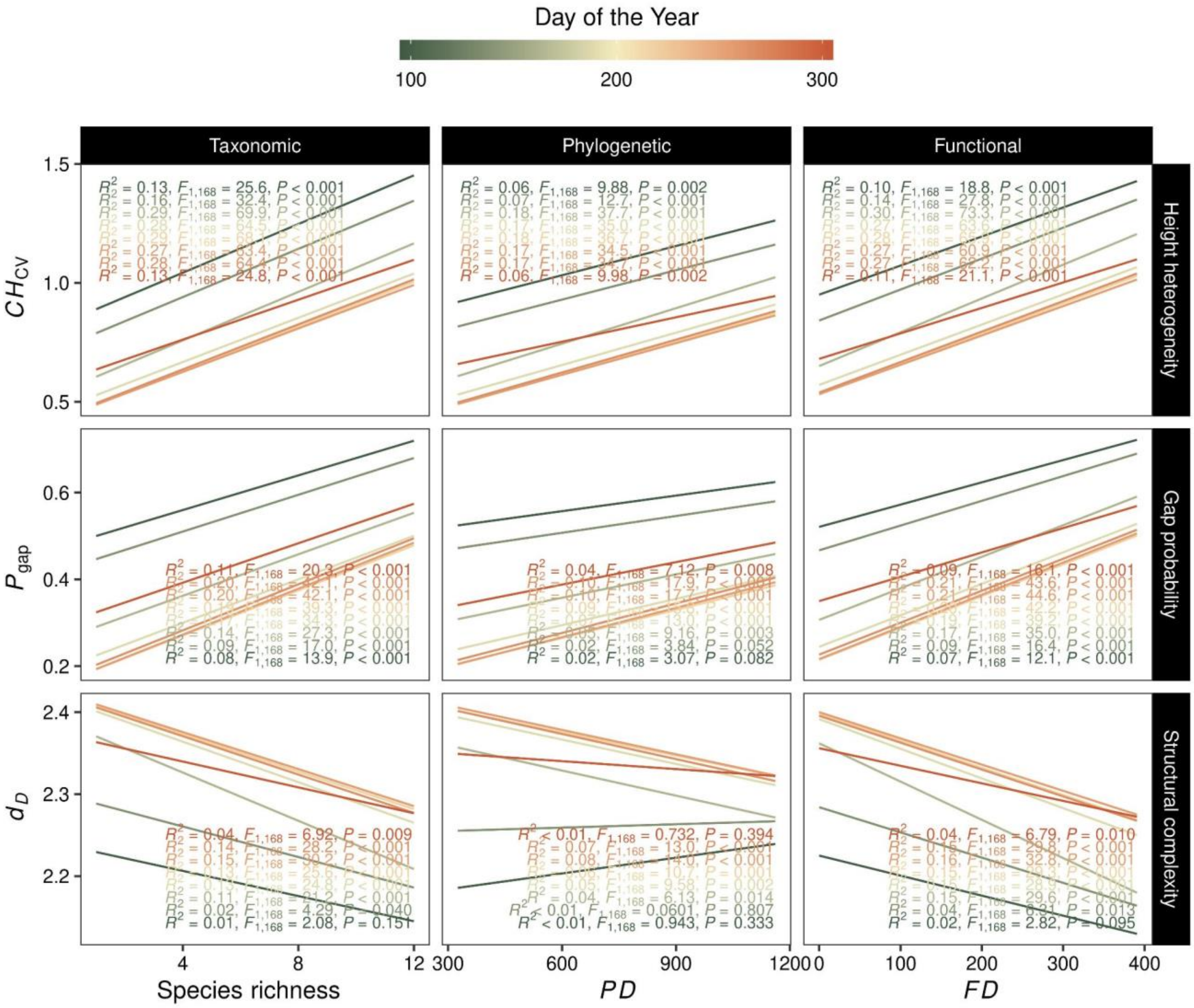


**Figure S6.** Influence of multiple dimensions of diversity on LiDAR-derived metrics at different observation periods during the growing season. $CH_{CV}$ describes the coefficient of variation of canopy height, $P_{gap}$ the gap probability, and $d_D$ the fractal dimension.

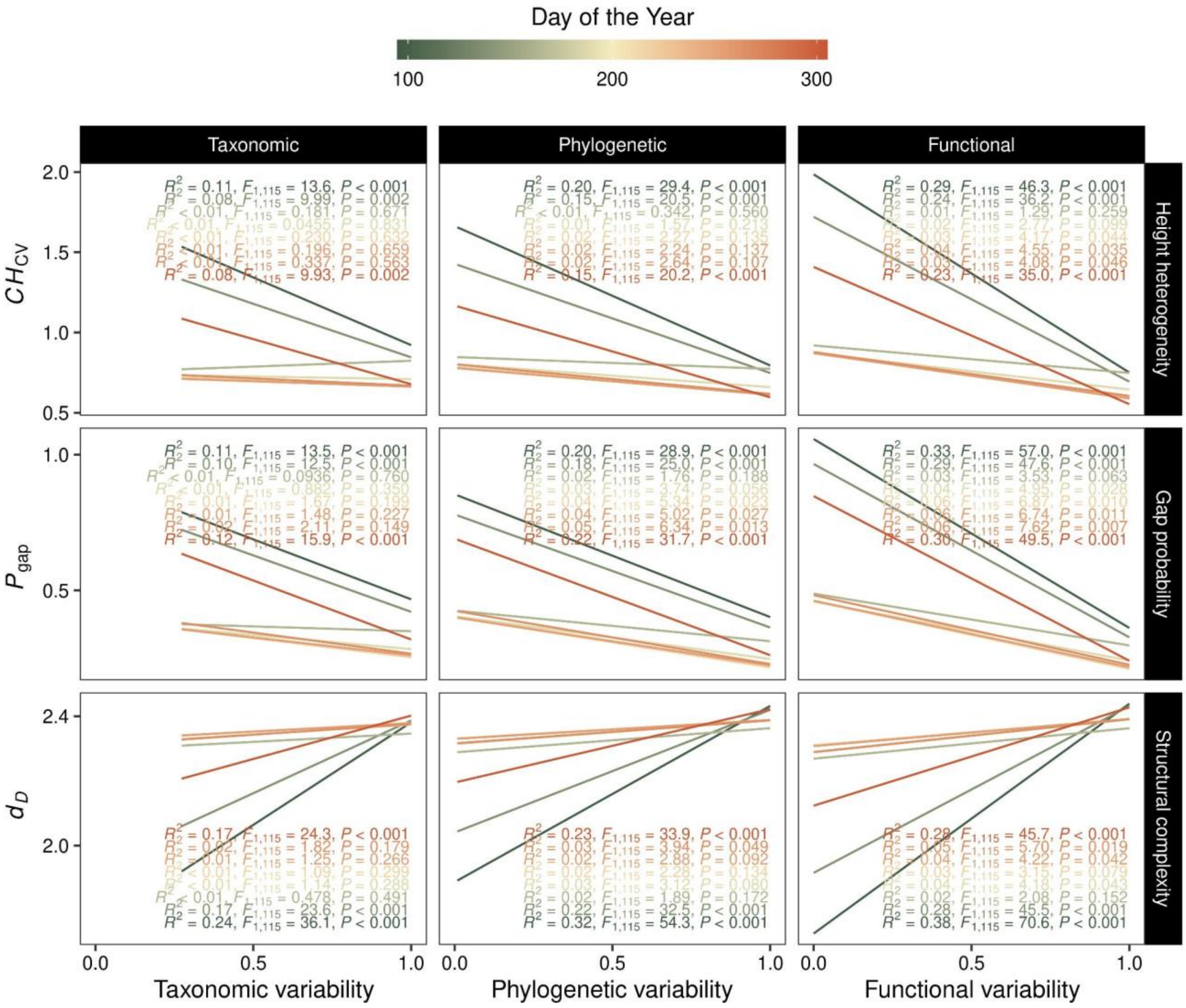


**Figure S7.** Influence of multiple dimensions of variability on LiDAR-derived metrics at different observation periods during the growing season. $CH_{CV}$ describes the coefficient of variation of canopy height, $P_{gap}$ the gap probability, and $d_D$ the fractal dimension.

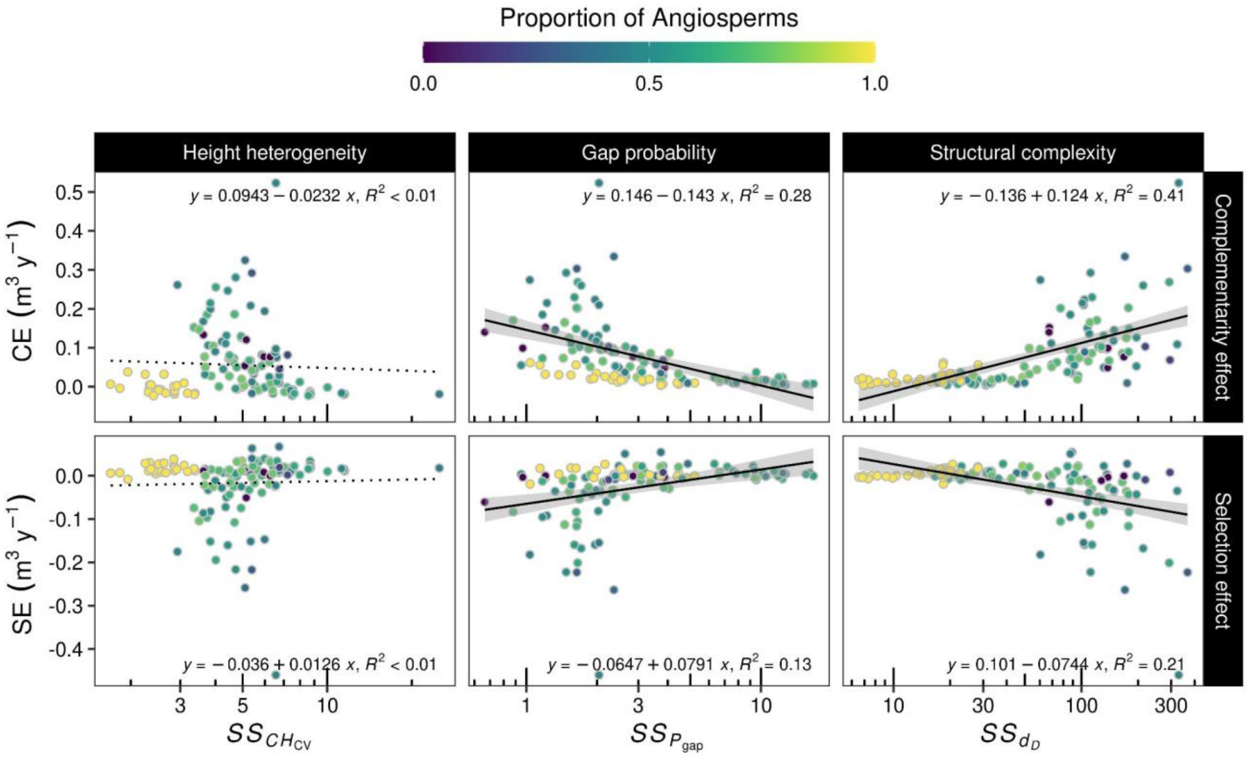


**Figure S8.** Effect of seasonal structural stability (*SS*) of LiDAR-derived metrics on complementarity (CE) and selection effect (SE) of annual wood productivity. $CH_{\mathrm{CV}}$ describes the coefficient of variation of the canopy height, $P_{\mathrm{gap}}$ the gap probability, and $d_{\mathrm{D}}$ the fractal dimension. Colors represent the proportion of angiosperms trees that were planted for each plot.

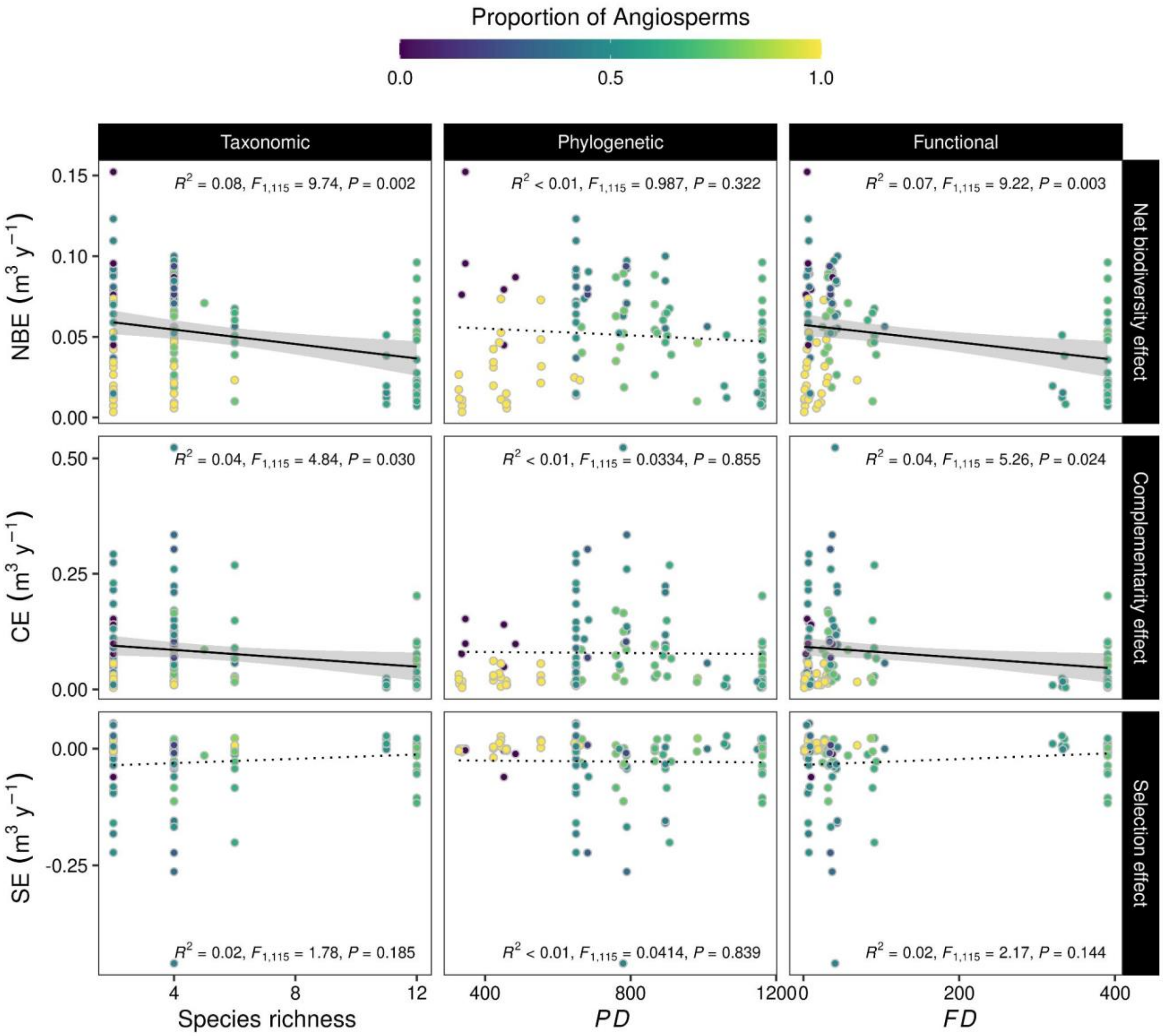


**Figure S9.** Influence of multiple dimensions of diversity on the net biodiversity (NBE), complementarity (CE), and selection (SE) effects of annual wood productivity.

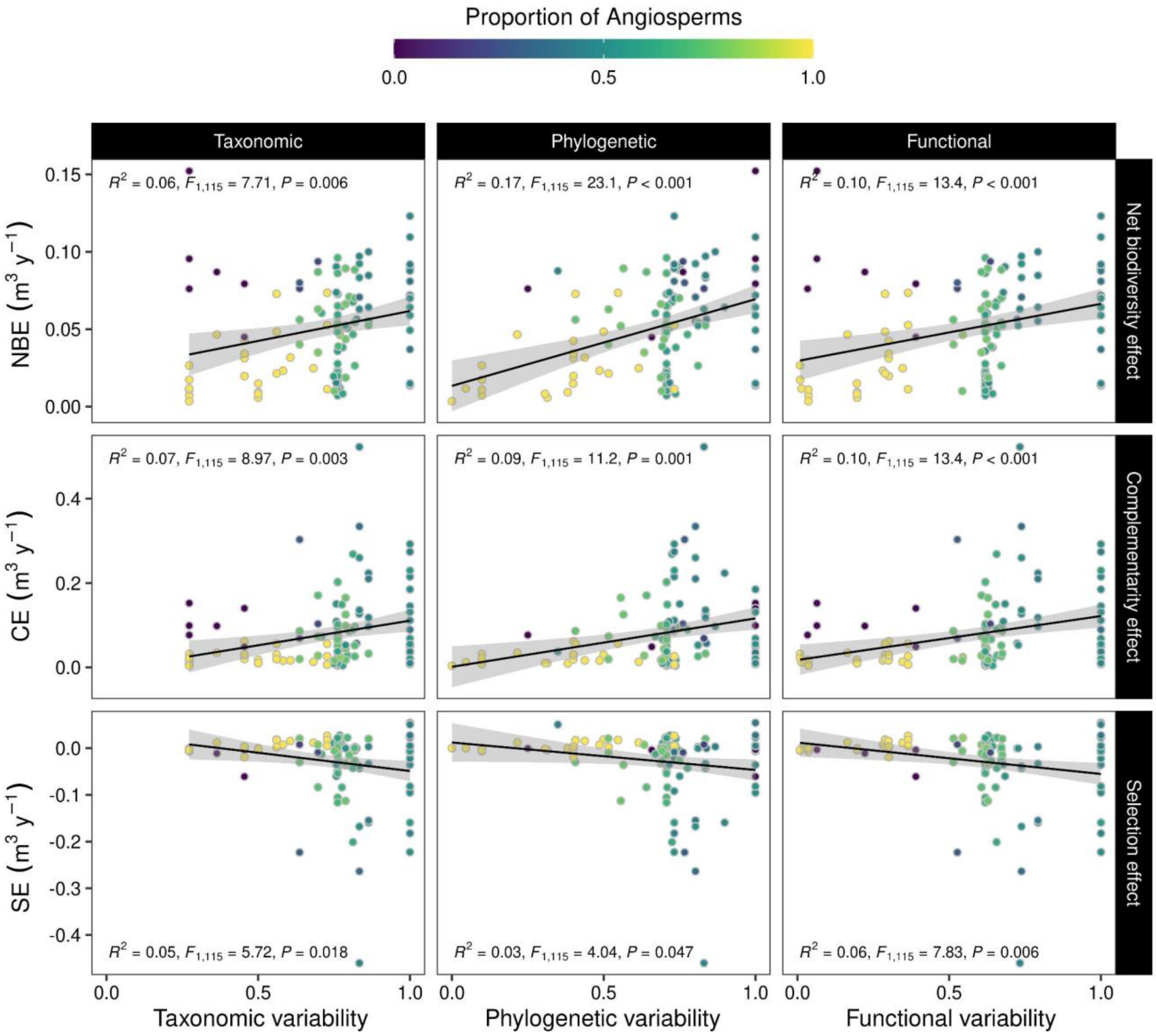


**Figure S10.** Influence of multiple dimensions of species variability on the net biodiversity (NBE), complementarity (CE), and selection (SE) effects of annual wood productivity.

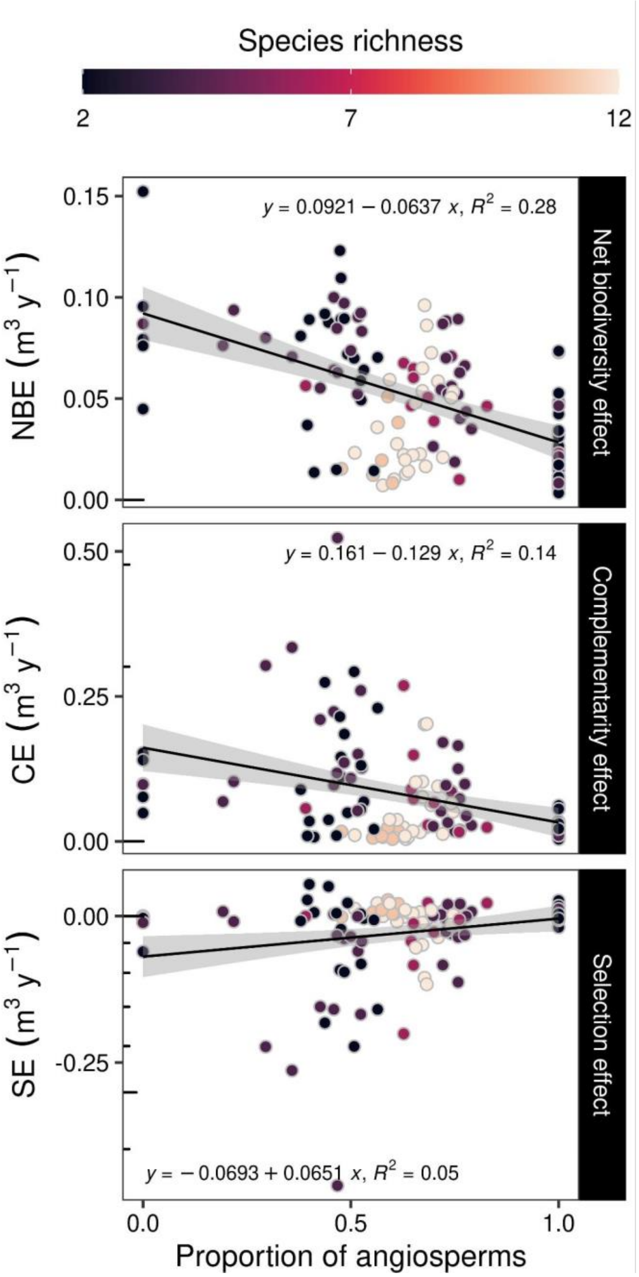


**Figure S11.** Effect of the proportion of angiosperm trees on the net biodiversity (NBE), complementarity (CE), and selection (SE) effects of annual wood productivity. Each point represents a plot.

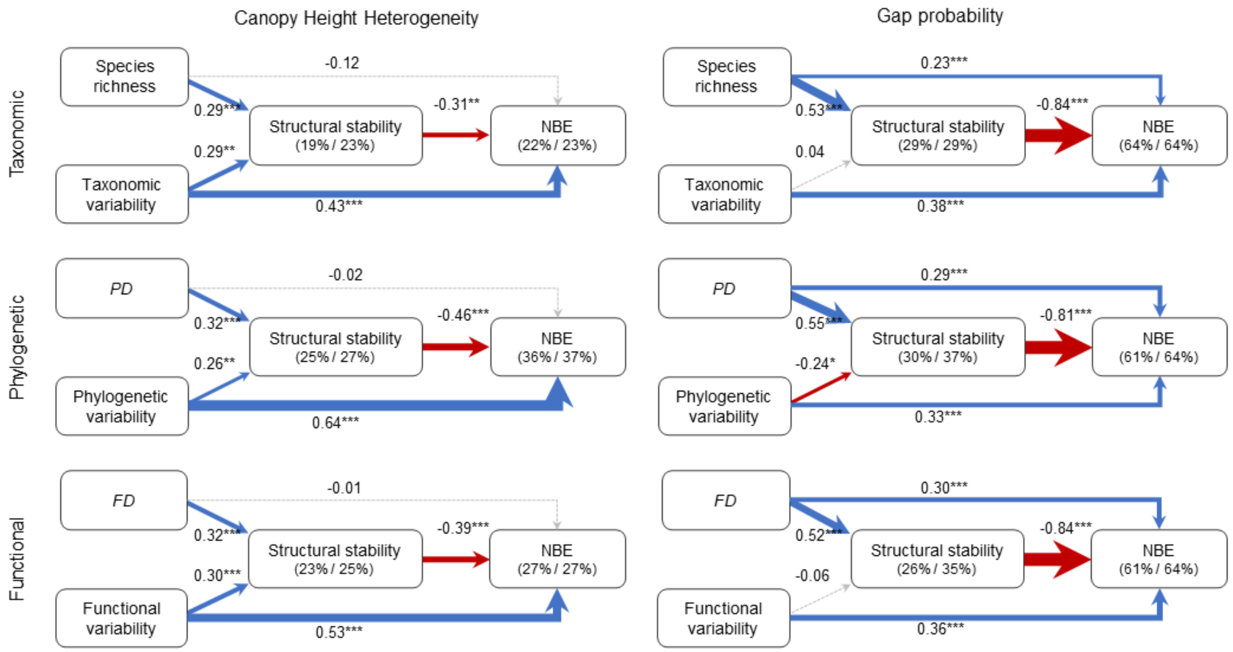


**Figure S12.** Structural equation models to describe paths of how multiple dimensions of diversity and variability influence seasonal structural stability of the canopy height heterogeneity and gap probability across the growing season, and thus, the net biodiversity effect (NBE) on productivity (i.e., overyielding). Values next to the arrows represent the standardized coefficients and their significance, while values in parentheses show the coefficient of determination of marginal and conditional effects. The thickness of the arrows indicates the standardized coefficients, while their color is positive (blue), negative (red), or non-effect (grey).

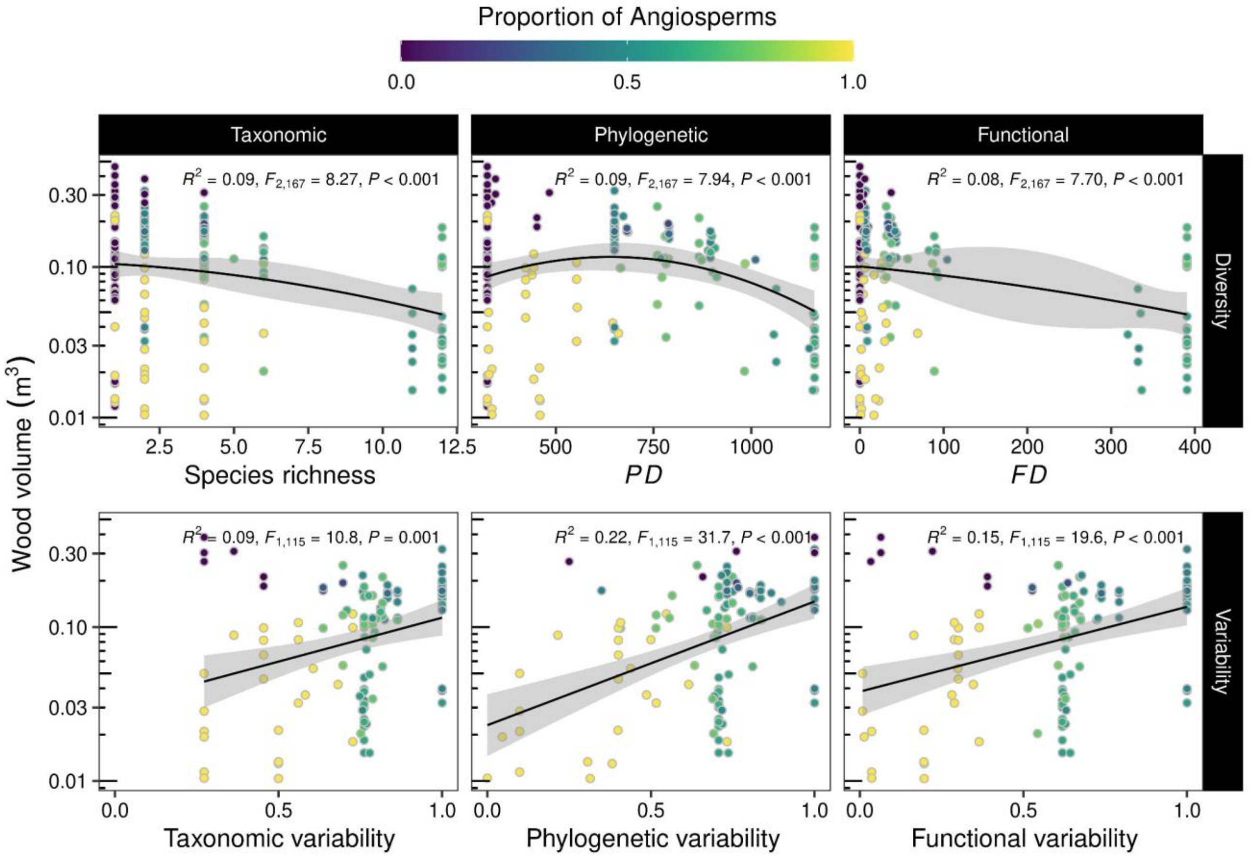


**Figure 13.** Influence of multiple dimensions of diversity and variability on wood volume.
